# A Framework for Benchmarking Pathway Reconstruction Algorithms

**DOI:** 10.64898/2026.08.04.742550

**Authors:** Neha Talluri, Tristan Figueroa-Reid, Justin Hiemstra, Chris S. Magnano, Adam Shedivy, Nistha Panda, Yancheng Liu, Sumedha Sanjeev, Oliver Faulkner Anderson, Altaf Barelvi, Aden O’Brien, Olivia T. Johnson, James A. Haddad, Spencer A. Halberg-Spencer, Akniyet Nurbol, Iris Jan, Mahunan Degbelo, Daniel Nachreiner, Canek Llera-Magord, Gabriel Howland, Grace H. Li, Anna Ritz, Anthony Gitter

**Author notes:** Department of Computer Science, Tufts University, Medford, MA, USA. Department of Computer Science, Brown University, Providence, RI, USA. Department of Computer Science, University of Colorado Boulder, Boulder, CO, USA. These authors contributed equally to this work (Anna Ritz, Anthony Gitter).

## Abstract

Cells coordinate diverse biological processes through interactions among thousands of molecules, but mapping these interactions comprehensively and systematically remains an open problem. Pathway reconstruction algorithms address this problem by linking molecules of interest, identified from high-throughput omics experiments, using prior knowledge encoded as background interaction networks. This process recovers intermediate molecules and interactions that were not directly measured in the experimental data but plausibly connect the observed molecules. It generates testable hypotheses about interactions that drive cell behavior and informs the choice of follow-up experiments. Many algorithms have been created over decades, each optimizing different computational objectives and relying on different assumptions. The resulting heterogeneity has made benchmarking challenging, limiting systematic comparisons. Therefore, selecting an algorithm for a given biological context remains a non-trivial and poorly informed task. This registered report presents a large-scale benchmark of pathway reconstruction algorithms, evaluating 14 algorithms across 822 datasets from four biological settings. To enable this benchmark, we introduce Signaling Pathway Reconstruction Analysis Streamliner (SPRAS), which standardizes algorithm inputs, outputs, and execution into a formal framework, enabling systematic comparison that was previously infeasible. We will assess each algorithm on reconstruction performance against gold standard pathways, algorithm similarity, and computational performance across different biological contexts. Together, these evaluations will provide quantitative evidence for understanding pathway reconstruction algorithm behavior and guiding algorithm selection.

## 1 Introduction

Understanding how molecules (e.g., proteins, RNA, DNA) interact in a cell to orchestrate biological processes is an open challenge. High-throughput omics technologies (e.g., transcriptomics, proteomics) allow us to measure thousands of molecular features of a biological system, providing a snapshot of how cells respond to disease, perturbations, and environmental conditions. These technologies identify which molecules are active in a given condition, but do not reveal which interactions among them are actually driving that condition.^1, 2^ Molecular interaction networks provide a model for addressing this challenge, organizing omics measurements into models of cellular behavior that capture which molecules are active, how they interact, and what functional roles they play.^2, 3^

For decades, the need for these molecular networks has motivated the development of pathway databases, curated repositories that compile experimental and literature-derived interactions into network representations of metabolic, signaling, and regulatory processes.^4^ Databases such as Reactome,^5^ KEGG,^6^ and PANTHER^7^ established the foundation for representing pathways computationally, enabling analyses like network pattern detection, comparison with expression data, and simulations that were previously infeasible.^4^ However, these databases represent consensus knowledge aggregated across many conditions, experiments, and cell types. They do not capture the interaction patterns that arise under specific experimental conditions.^8^ For example, an interaction present in a canonical pathway may be active in healthy tissue but absent or rewired in a disease state, so a database’s consensus representation can misrepresent the pathway actually operating in a given condition.

Pathway reconstruction algorithms address this gap by integrating context-specific omics measurements with a prior knowledge background interaction network, called an interactome, to infer condition-specific pathways that effectively represent hypotheses about which molecules are active and the nature of their interactions.^9^ Given a background interactome and a set of nodes of interest, an algorithm selects a pathway connecting those nodes by adding intermediate nodes and edges from the interactome as needed, balancing recovery of the nodes of interest against the number of new nodes and edges added.

The lack of independent benchmarks and evaluation datasets to assess algorithmic assumptions is one of the main open challenges for pathway reconstruction.^1^ Pathway reconstruction encompasses a large and heterogeneous space of algorithm formulations, spanning methodological classifications including network flow, Steiner trees, edge orientation, random walks, and disease module identification. When applied to the same dataset, these algorithms can yield entirely different pathways, making algorithm selection for a specific biological context non-trivial. Few comparative studies exist to guide this selection in large part because the diverse software dependencies, data formats, and parameter spaces of these algorithms create substantial technical barriers for systematic benchmarking at scale. As a result, algorithm selection defaults to familiarity, community popularity, or perceived fit with a dataset rather than demonstrated effectiveness for specific biological contexts.

We will benchmark pathway reconstruction algorithms at scale, evaluating 14 algorithms across 822 datasets from four pathway reconstruction applications (Figure 1). Our Signaling Pathway Reconstruction Analysis Streamliner (SPRAS) software unifies pathway reconstruction algorithms under a shared framework that standardizes inputs, outputs, and execution into a formal workflow, enabling systematic benchmarking that was previously infeasible and supporting best practices for computational bench-marking.^10^ SPRAS includes algorithms not typically categorized as pathway reconstruction methods, broadening the scope of comparison. We will assess each algorithm across all datasets on reconstruction performance against gold standard data, algorithm similarity, and computational performance. We will also introduce an algorithm-agnostic parameter tuning procedure that enables systematic parameter selection without method-specific assumptions. Together, these assessments will provide evidence for understanding pathway reconstruction algorithm behavior and guiding algorithm selection across biological contexts.

**Figure 1.**
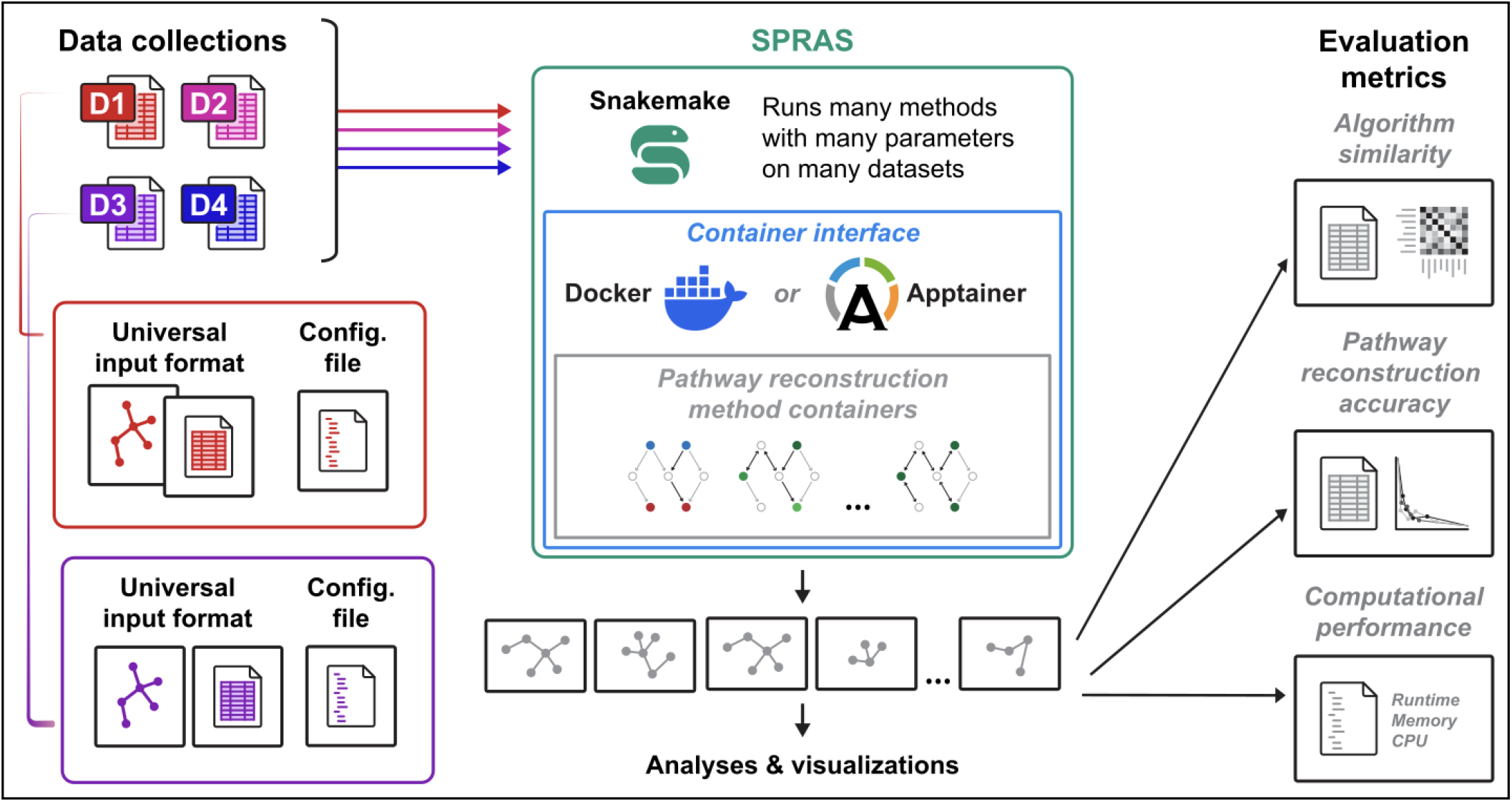
Overview of our benchmarking framework. Multiple data collections (D1-D4), representing different pathway reconstruction applications, will be converted to a universal input format each with an accompanying configuration file. Each of these will be passed to SPRAS, which uses Snakemake^11^ to run multiple pathway reconstruction algorithms across multiple parameter settings. The resulting reconstructed pathways will be evaluated across three assessments: algorithm similarity, pathway recovery performance, and computational performance.

## 2 Background

Below, we provide core concepts on pathway reconstruction algorithms, their data inputs, and their output representations to establish the terminology and concepts used throughout this paper.

### 2.1 Pathway Reconstruction Algorithms

Pathway reconstruction algorithms are a broad group of network biology methods that aim to identify a subset of nodes and edges specific to a pathway or condition from a larger network. They define an objective function over two inputs: a prior knowledge background interaction network (interactome) and context-specific omics measurements (input nodes). The algorithms apply the optimization function to select a reconstructed pathway of nodes and edges from the interactome that connects the input nodes and represents a process or cellular state under a condition of interest. The resulting pathway balances the inclusion of the input nodes against the number of additional nodes and edges drawn from the interactome. This task has also been referred to as network inference,^12^ pathway inference,^3^ network reconstruction,^3, 13^ subnetwork reconstruction,^13^ subnetwork inference,^13^ and signaling network reconstruction.^3^

Pathway reconstruction algorithms span several methodological classifications. Steiner forest algorithms address a network design problem where the goal is to establish connections among special vertices designated as terminals at minimum cost.^14^ These algorithms connect terminal nodes (e.g. differentially expressed molecules) through an interactome by finding minimum-cost trees or forests that span these terminals, often incorporating intermediate non-terminal nodes (Steiner nodes) to explain signal propagation. Flow-based algorithms model propagation as network flow, where edge capacities or costs reflect interaction confidence, activity levels, or weights. These algorithms find pathways that maximize flow from source nodes to target nodes, tracing how biological signals move through an interactome. Edge orientation algorithms assign edge directions in an undirected interactome by optimizing a connectivity objective.^15^ Network propagation algorithms diffuse information across connected nodes in a network.^16^ This class assumes that nodes near one another, or connected through shared neighbors, tend to share function or phenotype, so signal from a set of important nodes is spread through the interactome to prioritize every node by its proximity to them.^16^ Random walk-based algorithms are a subset of this class: they explore the global topology of networks by simulating a walk that iteratively transitions from a node to one of its neighbors.^17, 18^

Other types of graph algorithms also fit this definition without being explicitly designed for pathway reconstruction. Disease module identification methods (also referred to as active module identification methods^19^ or module detection^20^) identify densely connected communities within an interactome that correspond to disease-associated pathways and mechanisms,^21^ rather than selecting edges to connect a specified set of nodes. The outputs from disease module identification methods are comparable with pathway reconstruction outputs, so we include them in our benchmarking to evaluate them on pathway reconstruction tasks.

### 2.2 Input Data

Pathway reconstruction algorithms require at minimum two inputs: an interactome and a set of nodes. The interactome encodes prior knowledge of molecular relationships as a graph, where nodes represent molecules and edges represent experimental, literature-supported, or predicted interactions between them. Input nodes are a subset of nodes in the graph corresponding to molecules of interest, derived from high-throughput omics data, manual curation, or literature. The input nodes are mapped onto the interactome, and algorithms build pathways from the interactions that involve those input nodes.

The background interactome serves as prior knowledge of molecular interactions among a defined set of biological entities. Publicly available interactomes are constructed from diverse biochemical and omics data sources, representing literature-curated and experimentally-measured interactions among biological entities.^22^ These interactions span protein-protein interactions, regulatory relationships, signaling cascades, metabolic reactions, and functional associations.^22^ The interactome can also encode the direction of interactions. Edges can be directed, where one molecule regulates or controls another, or undirected, such as membership in a complex or co-participation in an interaction where no causal direction is implied.^23^ Interactomes can represent directionality explicitly, yielding fully directed or mixed interactomes (containing both directed and undirected edges), or omit it entirely, yielding fully undirected interactomes where all edges are treated as bidirectional. Pathway reconstruction algorithms vary accordingly, accepting fully directed, fully undirected, or mixed interactomes depending on their underlying methods.

Input nodes may take different forms depending on each algorithm’s requirements. Source and target nodes define the start and end points of the pathway. Sources represent upstream initiating molecules such as ligands, receptors, or perturbed proteins. Targets represent downstream effectors such as transcription factors (TFs) or differentially expressed genes. Algorithms using this format find pathways connecting sources to targets through the interactome. Other algorithms take a single set of input nodes with numerical scores called prizes. Prizes represent the nodes’ relevance to a biological condition and may be derived from omics data such as phosphoproteomics, mutation profiles, differential expression, or genome-wide association study (GWAS) hits, where higher values indicate greater evidence of relevance. Active nodes (also known as seeds^19^) encode a binary version of prizes, marking a node as relevant. Algorithms using these formats find pathways that maximize the inclusion of prize nodes or active nodes. Dummy nodes are artificial root nodes that give the algorithm a starting point (analogous to a source node).^24^

### 2.3 Output Pathways

Pathway reconstruction algorithms output a reconstructed pathway, a subset of nodes and edges from the background interactome. The output represents a candidate pathway, that is, a hypothesis about the observed biological interactions and active molecules. Reconstructed pathways can be undirected, directed, or mixed in edge directionality depending on the algorithm interpretation and the directionality of the input interactome. Some algorithms assign weights or scores to output nodes and edges reflecting their rank or confidence in the reconstructed pathway, while others return unweighted pathways.

## 3 Methods

This section describes the methods we will use to benchmark pathway reconstruction algorithms. We first describe the 14 algorithms selected for benchmarking and the four dataset collections that will be used to evaluate them. We then describe the evaluation metrics that will be used to assess each algorithm, followed by the parameter tuning procedure that will be used to select algorithm parameters for each dataset. We next describe SPRAS, the framework developed to standardize algorithm inputs, outputs, and execution. We describe how we will use SPRAS to evaluate each algorithm’s pathway recovery performance, algorithm similarity, and computational performance across all dataset collections. Finally, we describe the infrastructure we will use to run the benchmark at scale.

### 3.1 Benchmarked Pathway Reconstruction Algorithms

We will benchmark 14 pathway reconstruction algorithms that were chosen to represent a wide range of methodological classifications (Figure 2).

**Figure 2.**
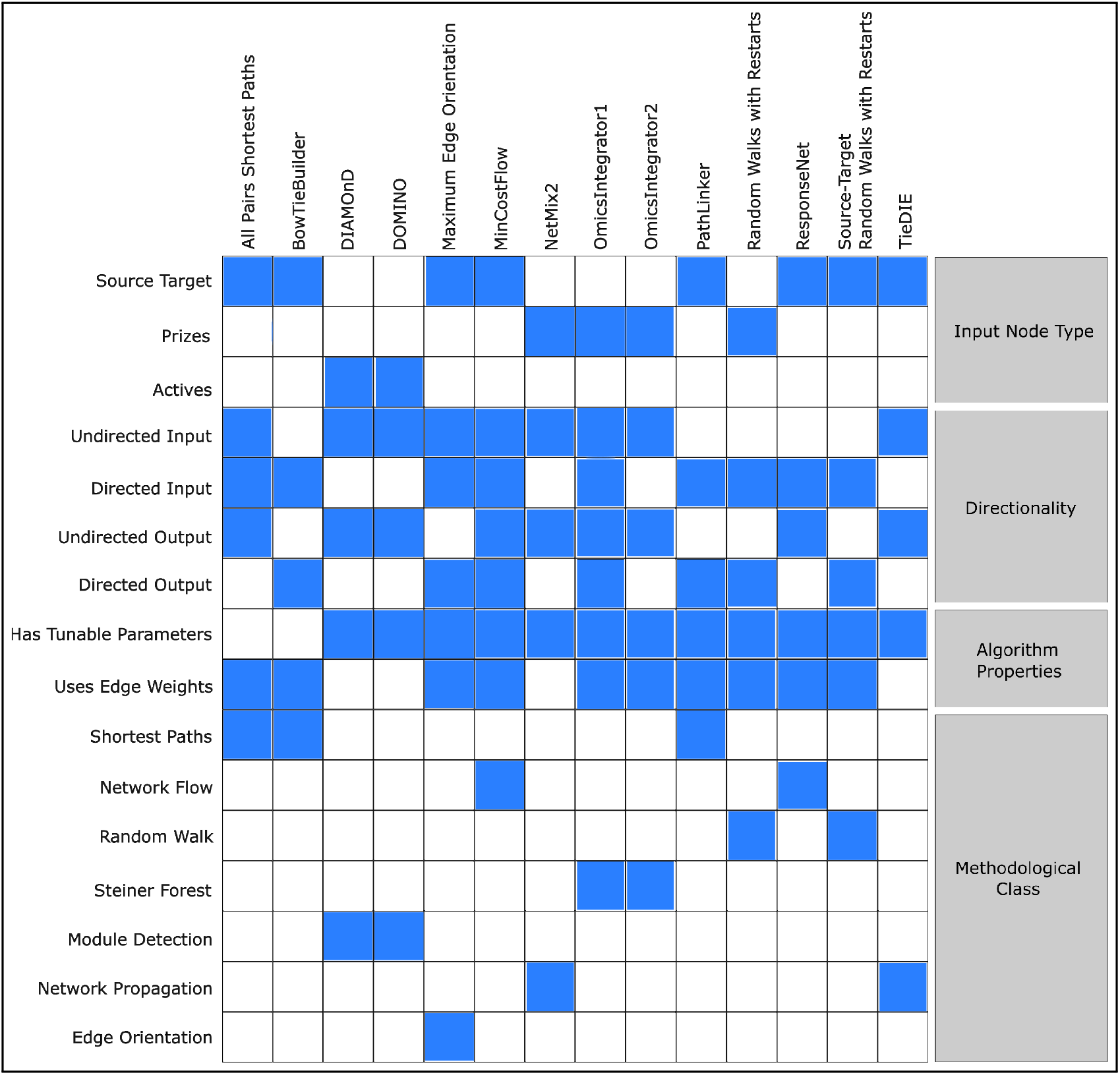
Overview of pathway reconstruction algorithm features. Rows correspond to algorithm features grouped into four categories. Filled cells indicate that an algorithm has the corresponding feature. Further details are provided in Supplementary Table 1.

**All Pairs Shortest Paths (APSP)**^25^ reconstructs pathways by computing the shortest path between every source-target pair and returns the union of these paths as the reconstructed pathway.

**BowTieBuilder (BTB)**^26^ reconstructs pathways by iteratively connecting sources and targets to a growing pathway. At each iteration, it finds the shortest path from any input node to the existing pathway and adds it to the pathway, repeating the process until all sources and targets are connected or no more connections are possible.

**DIAMOnD**^20^ reconstructs pathways by greedily selecting high-connectivity significant nodes near active nodes and ranking them by their connectivity p-values, returning the induced pathway generated by a user-specified number of the highest-ranking nodes.

**DOMINO**^27^ reconstructs pathways by taking several steps to find highly connected pathways surrounding active nodes, then identifying and returning pathways with active node overrepresentation.

**Maximum Edge Orientation (MEO)**^28^ reconstructs pathways by orienting edges in networks to maximize connectivity from source to target nodes by assigning directions to undirected edges to create high-confidence pathways that maximize the total edge weight.

**MinCostFlow (MCF)**^29–31^ reconstructs pathways by formulating pathway reconstruction as a minimum-cost flow problem identifying paths that connect sources to targets. Our implementation of MCF is inspired by ResponseNet.^32^

**NetMix2**^33^ reconstructs ‘altered pathways’, or what we call disease modules, by first estimating the number of nodes *n* in the candidate altered pathway of the input seed nodes, then solving an optimization problem to return a connected pathway of size *n* that maximizes the input node scores.

**Omics Integrator 1 (OI1)**^24^ reconstructs pathways using the prize-collecting Steiner forest (PCSF) formulation,^34^ which finds pathways that maximizes node prizes and minimizes edge costs. Prizes are assigned to terminal nodes (e.g., proteins identified from omics data), and edge costs reflect interaction confidence in the underlying interactome. The optimization problem is solved with the msgsteiner message-passing algorithm.^35^

**Omics Integrator 2 (OI2)**^36^ is an updated version of Omics Integrator 1 that solves the PCSF formulation using the pcst_fast solver^37^ in place of msgsteiner.

**PathLinker**^38, 39^ reconstructs pathways by finding the *k* highest scoring paths between sources and targets. The algorithm integrates Yen’s algorithm with an A* heuristic, ranking molecules and interactions based on their first occurrence across *k* shortest paths.

**Random Walks with Restarts (RWR)**^18^ reconstructs pathways by scoring every node by its proximity to the prize nodes. At each step the walker moves to a neighbor or, with fixed restart probability, teleports back to the prize nodes. The algorithm natively returns a set of nodes, not a pathway. To produce a pathway, we retain the nodes with output scores above a threshold and then induce a subgraph on the interactome restricted to edges where both endpoints fall within this node set.

**ResponseNet**^32^ reconstructs pathways by solving a minimum-cost flow optimization problem using linear programming, balancing the cost of adding edges against the flow through the network to identify high-confidence connections between sources and targets.

**Source-Target Random Walks with Restarts (ST-RWR)**^18^ reconstructs pathways by running random walks with restarts twice: once from the source nodes and again from the target nodes (with directed edges reversed). ST-RWR identifies nodes ranked highly under both walks. Similar to RWR, we retain the nodes with scores above a threshold and induce a subgraph on the interactome. Our implementation of ST-RWR is inspired by TieDIE.^40^

**TieDIE**^40^ reconstructs pathways using a diffusion method that connects sources and targets through linkers found using forward diffusion from the sources and backward diffusion from the targets, then constructs a pathway connecting sources to targets through the linkers.

### 3.2 Dataset Collections

We will evaluate the 14 algorithms across four dataset collections. A collection is a group of related dataset instances that generated data in the same way but span different biological contexts. These collections range from curated pathway-derived datasets, which offer controlled settings with an expectation of how the pathway should be structured, to omics-derived datasets, which reflect the complexity of real experimental measurements.

A dataset is a single instance within a dataset collection (e.g., one cell line within a cancer cell line collection) defined by a distinct set of inputs (input nodes and interactome) and a set of nodes and/or edges to evaluate the reconstructed pathways. We refer to this set of nodes and/or edges as the ‘gold standard’ for that input because it will be used to calculate evaluation metrics. However, this term is used more loosely in this benchmark than in other biological contexts. In some biological settings, human annotations can provide the gold standard, such as in microscopy image segmentation.^41^ In others, a biological assay may provide a strong signal to use as the gold standard, such as ChIP-exo^42^ for predicting TF-DNA binding along a genome.

Biological pathways lack a perfect gold standard. Some of our datasets come from pathway databases that characterize nodes and edges in each pathway. Here the modeling task is to recover parts of the pathway given some of the nodes as input. The pathway’s nodes and edges serve as the gold standard. In other cases, the input data comes from omics assays, and the complete underlying biological process is unknown. In these datasets, we use complementary biological data to assess the overlap between the nodes in the predicted pathway and the nodes highlighted by the complementary data. Strong node overlap increases our confidence in the predicted pathways, but it cannot quantify edge-level predictions and does not reflect the true unknown nodes involved in the pathway.

For all datasets, we restrict sources, targets, and gold standard data to only include molecules and interactions present in the background interactome. This ensures that algorithm performance will reflect the methods themselves rather than incompleteness in the background network. Without this filtering, an algorithm’s inability to recover a gold standard edge could reflect either algorithmic limitations or the absence of that edge from the interactome; likewise for nodes.

#### Input Background Interactome

The main background interactome used for all dataset collections is the human STRING v12 physical interactome.^43^ Pathway reconstruction algorithms operate on the assumption that edges represent direct molecular interactions between proteins, which motivates the choice of the physical network over the functional network. The physical network indicates whether two proteins are in physical proximity, for example through direct binding or membership in the same molecular complex.^43^ The original interactome has 738,805 undirected edges. Each edge carries a combined score derived from the subset of STRING evidence channels that support physical interactions: text mining, experiments, and curated databases.^43^ The scores range from 150 to 999, with larger values indicating higher confidence.^43^ The score reflects how strongly the available data support the presence of a biological association.^43^ We threshold the interactome at a score of ≥ 400 (the medium confidence threshold), leaving 210,667 undirected edges.^43^

##### 3.2.1 PANTHER Signaling Pathways

This dataset collection contains curated human signaling pathways describing molecular interactions underlying cell signaling processes. A signaling pathway is a series of molecular interactions within a cell, initiated by ligand-receptor binding or other stimuli, propagated through intermediate molecules, and ending in specific changes to gene expression, protein activity, or cell behavior.^44^ We use a library of curator-defined protein pathways and groupings called PANTHER Pathways^7^ present in Pathway Commons.^45^ We restrict this dataset collection specifically to signaling pathways. The goal of this dataset collection is to test whether a pathway reconstruction algorithm can recover the biological knowledge already encoded in a curated pathway. This is best described as controlled pathway recovery. The algorithms reconstruct structure that is already known rather than predicting entirely novel interactions.

##### Chosen PANTHER Pathways

PANTHER does not formally classify signaling pathways,^7^ so we defined our own criteria. We downloaded the list of *Homo sapiens* PANTHER pathways, which contains 159 pathways. We partitioned these into four groups based on specific terms in their pathway names: 46 signaling pathways (pathway names containing the word ‘signaling’ or ‘signalling’), 12 metabolic pathways (pathway names containing ‘metabolic’), two hormone pathways (pathway names containing ‘hormone’), and 99 other pathways.

We then input the 99 remaining pathway names into Claude Opus 4.6^46^ with the prompt: ‘From this text file that is full of pathway names, can you take everything that is potentially a cell signaling pathway and rank them by the likelihood they are a signaling pathway? None of the pathways provided will have the word signaling in the names.’ Claude proposed 38 candidate signaling pathways from this set. We then manually reviewed these 38 pathways. We also reviewed the 46 original signaling pathways to verify each was a *Homo sapiens* pathway. Despite filtering by species during download, PANTHER returned some non-human pathways.

We were left with 29 candidate signaling PANTHER pathways (20 chosen from the 46 signaling pathways from PANTHER and 9 chosen from the 38 Claude proposed pathways). We then filtered for pathways present in Pathway Commons and for pathways with at least one source node and one target node. Pathways lacking sources or targets cannot be used with algorithms that require source-target pairs, so we excluded them to construct a dataset compatible with all algorithms used in this benchmark. After filtering, 22 PANTHER pathways remained (Supplementary Table 2).

In addition to the 22 PANTHER pathways, we constructed several merged pathways. The first merged pathway, mega pathway, combines all 22 pathways into a single pathway and input node set. We also constructed target-focused and source-focused merged pathways. The target-focused version merged all pathways with more than 10 targets (excluding Wnt, which also has 106 sources and would dominate the source side), and the source-focused version merged all pathways with more than 10 sources (excluding Cadherin and Wnt, which each contribute disproportionately to the source side). Finally, we constructed a balanced merged pathway using a greedy expansion procedure: starting from the single most balanced pathway (closest to a 1:1 source to target ratio), we iteratively added pathways that maximized merged node count while keeping the source/target imbalance within a tolerance of 0.15, stopping when no remaining pathway could satisfy the tolerance constraint. A full list of the merged pathways is provided in Supplementary Table 3.

##### Input Nodes

The sources for this dataset are cell surface receptors. We use predicted human cell-surface receptor proteins generated by SURFY,^47^ a random forest classifier trained on experimentally-verified cell-surface proteins from the Cell Surface Protein Atlas.^48^ SURFY predicted 2, 886 surface-exposed proteins across human cell types and developmental stages.^47^ The targets for this dataset are human TFs. For the human TFs, we use the Animal Transcription Factor Database (AnimalTFDB) v4.0.^49^ AnimalTFDB 4.0 identifies 1,659 TFs.

For each PANTHER pathway, a protein is assigned as a source if it appears in both the pathway and the human receptor dataset or as a target if it appears in both the pathway and the human TF dataset. Prize and active nodes are both defined as the union of source and target nodes. Prizes are assigned a uniform score of 1.0 reflecting binary pathway membership. Each PANTHER pathway yields a distinct set of input nodes comprising sources, targets, prizes, and actives.

##### Gold Standar

The gold standard for this dataset uses the full PANTHER pathways, with separate node and edge gold standards constructed for each pathway. The input nodes are, by construction, in the node gold standard, since both are derived from the same PANTHER pathway. The PANTHER pathways represent the current state of established biological knowledge, encoding interactions that have been validated and accepted by the community, making them a natural gold standard for evaluating whether a pathway reconstruction algorithm recovers known biology. These gold standards are also the only edge gold standards across all dataset collections; the other dataset collections lack edge level resolution.

##### Additional Input Background Interactomes

We merge the edges from all 22 chosen PANTHER pathways into the thresholded human STRING v12 physical interactome. When the pathways are combined with the interactome, this introduces directed edges as defined by Pathway Commons.^23, 45^ Those added pathway edges are given the median interactome edge weight.

In addition, we construct downsampled interactomes of varying sizes derived from the full physical interactome (without thresholding). The background interactome is downsampled by sampling a fixed fraction of edges in a stratified manner without replacement. Edges are stratified into bins by weight so that sampling preserves the overall STRING score distribution from the full physical STRING interactome. The downsampled interactomes are constructed at different increments [10%, 30%, 50%, 70%, 100%], with the corresponding fraction of edges drawn from each bin at each percentage.

For each PANTHER pathway, all pathway edges are concatenated with the downsampled interactome and those pathway edges are given the median interactome edge weight. Each resulting interactome is then evaluated against three criteria: 100% of targets must be reachable from at least one source, 100% of sources must be able to reach at least one target, and at least 30% of pathway edges must overlap with the sampled interactome. If the criteria are not met, sampling is repeated with a new random draw up to 50 attempts. If no attempt satisfies the criteria, the attempt with the highest pathway-interactome overlap is retained. This process yields five downsampled interactomes per pathway. Further information about each interactome is provided in Supplementary Tables 4 and 5.

When a PANTHER pathway is added to an interactome, its edges can duplicate edges already present in the interactome. To resolve these duplicates, we first prioritize the directed edge with the highest weight. If the duplicated edges are all undirected, we retain the edge with the highest weight.

#### 3.2.2 DepMap Cancer Cell Lines Collection

This dataset collection captures how upstream genomic alterations can propagate through signaling pathways to influence downstream gene expression within different cancer cell lines. We use data from DepMap 25Q3,^50, 51^ CCLE 2019,^52, 53^ and cBioPortal,^54–56^ which together provide omics measurements and molecular profiles across hundreds of cancer cell lines. From these data, we construct one SPRAS-formatted dataset per cell line, yielding 627 datasets in the collection, all run against a common background STRING interactome^43^.

##### Chosen Cell Lines

We restricted the cell lines to those present across all input datasets and the gold standard. After trimming the data to nodes in the interactome, we retained cell lines with at least one source node and one target node. This yielded 627 cell lines as the final set of cell lines in this collection (Supplementary File 1).

##### Input Nodes

The sources are derived from genetic events that are the primary causes for most cancers,^57^ somatic mutations and copy number alterations (CNAs). Both can dysregulate downstream signaling and propagate to changes in gene expression and protein activity. Somatic mutations are DNA changes acquired after conception, arising from replication errors during cell division and exposure to mutagens.^58^ CNAs occur due to changes to DNA structure that lead to the gain (amplification) or loss (deletion) of copies of DNA sections from a normal genome.^59^ Both can alter gene expression and protein activity; however, buffering mechanisms within a cell mean that not all CNAs produce proportional changes in altered gene or protein expression levels.^60^ Therefore, not all CNAs in a cell line are expected to have functional consequences.

The somatic mutations are from DepMap’s^50, 51^ damaging somatic mutations matrix,^a^ which encodes for each cell line whether each gene carries at least one likely loss-of-function variant, aligned to the h38 genome. The values follow a 0*/*1*/*2 scheme: 0 indicates no damaging mutation; for genes with one or more damaging mutations, allele frequencies are summed, and a value of 2 is assigned if the sum exceeds 0.95, otherwise 1 is assigned.^50, 51^ We use the scores of 1 or 2 for prizes because genes with a score of 2 have more importance than genes with a score of 1. For each cell line, genes with a score of 1 or 2 are retained as source input nodes in the cell line–specific dataset. Across cell lines, the number of somatic mutations ranges from 5 to 1,602.

The CNA data comes from CCLE and cBioPortal.^52–56^ Note we use the CCLE 2019 CNA release, which provides discrete amplification and deletion calls, rather than more recent DepMap releases, which only provide continuous copy number values. CCLE created a binary event matrix produced using the REVEALER method, which identified genomic alterations correlated with a phenotype, aligned to the hg19 genome.^61^ Each gene is represented by three binary features encoding mutation, amplification, and deletion status, where 1 indicates an alteration and 0 indicates none.^52, 53^ cBioPortal translated these binary calls to discrete CNA values, assigning amplifications a value of +2, deletions −2, and no event 0.^54–56^ The resulting values are restricted to {−2, 0, +2} and exclude the intermediate ±1 values present in the standard five-tier system, which distinguishes homozygous deletion, hemizygous deletion, neutral, gain, and high-level amplification.^62^ For each cell line, genes with a CNA value of −2 or +2 are retained as source input nodes. Across cell lines, the number of CNA genes ranges from 69 to 4,980. Each retained gene receives a prize of 0.5, a value below 1 to prevent CNA nodes from dominating reconstructed pathways relative to the number of somatic mutations and TFs, which are fewer in number per cell line.

The targets for this dataset are active TFs derived from gene expression data. To infer TFs that drive gene expression patterns per cell line, we use regularized network component analysis (NCA)^b^. NCA takes a gene expression matrix and a prior TF-target network and factorizes them into a matrix of TF activity (TFA) scores by samples and another of TF-to-target influence weights.^65^ A TFA score is a continuous value quantifying the inferred regulatory activity of a TF in a given cell line.

Unlike most of our other benchmarking datasets where a processed version of the inputs is already available, we compute the TF activities ourselves. Specifically, we use DepMap^50, 51^ bulk RNA-seq gene expression profiles,^c^ a cell line by gene matrix of log-transformed transcripts per million values from strand specific RNA-seq across 1,699 cell lines, aligned to the hg38 genome. Prior to running EstimateNCA, we merge duplicated cell lines by taking the mean of their expression values, z-score normalizing the expression values across cell lines, and removing genes expressed in fewer than 10% of cell lines. The prior TF-target network encodes putative regulatory relationships between TFs and their target genes.^63^ We use a hg38 motif-based regulatory prior ^d^ aligned to the same reference genome as the DepMap gene expression data. This prior was built by scanning for TF motif instances with motifmatchr^67^ and the CisBP reference motif database,^68^ linking each TF to a gene whose transcription start site lies within ±5kb of a matched motif instance.^63^

We run EstimateNCA with regularization parameter *λ* = 0.1, which controls sparsity and stabilizes TFA estimates. To assess variability in TFA scores, we generate 100 bootstrapped datasets by subsampling half of the cell lines with replacement (849 samples per bootstrap) and run EstimateNCA on each, producing 100 TFA profiles. To aggregate TFA profiles across the 100 bootstrap runs, we apply a sign-consistent averaging procedure to account for a known ambiguity in NCA: for any TF *i*, the observed target gene expression *e_i_* is equally explained by (*w_ai_, T_i_*) or (−*w_ai_,* −*T_i_*). *T_i_* is the TFA profile of TF *i*, and *w_ai_* are the regression weights learned between *T_i_* and *e_i_* for the nonzero elements of *a_i_*. Repeated runs may return TFA profiles that are identical up to a global sign flip. For each TF, we designate the activity vector from the first bootstrap as a reference and compute the Pearson correlation between it and the corresponding activity vector from each subsequent bootstrap, excluding missing values. Activity vectors with negative correlation are sign-flipped before addition; those with positive correlation are added directly. The running sum is divided elementwise by a per-TF count of contributing runs. All TFs appeared in all 100 bootstrap runs with the exception of two TFs, HOXC11 and SOX7, which appeared in 99 and 98 runs respectively. This yields a single consensus TFA matrix of TFs by cell lines representing the regulatory activity estimate for each TF.

After computing the consensus TFA matrix, we z-score normalize TFA scores across cell lines for each TF to place all TFs on a common scale (Figure 3). For each cell line, we retain TFs whose normalized activity meets or exceeds two standard deviations from the mean; cell lines with at least one qualifying TF are used. These TFs serve as target input nodes in each cell line-specific dataset. Across cell lines, the number of TFs ranges from 1 to 121.

**Figure 3.**
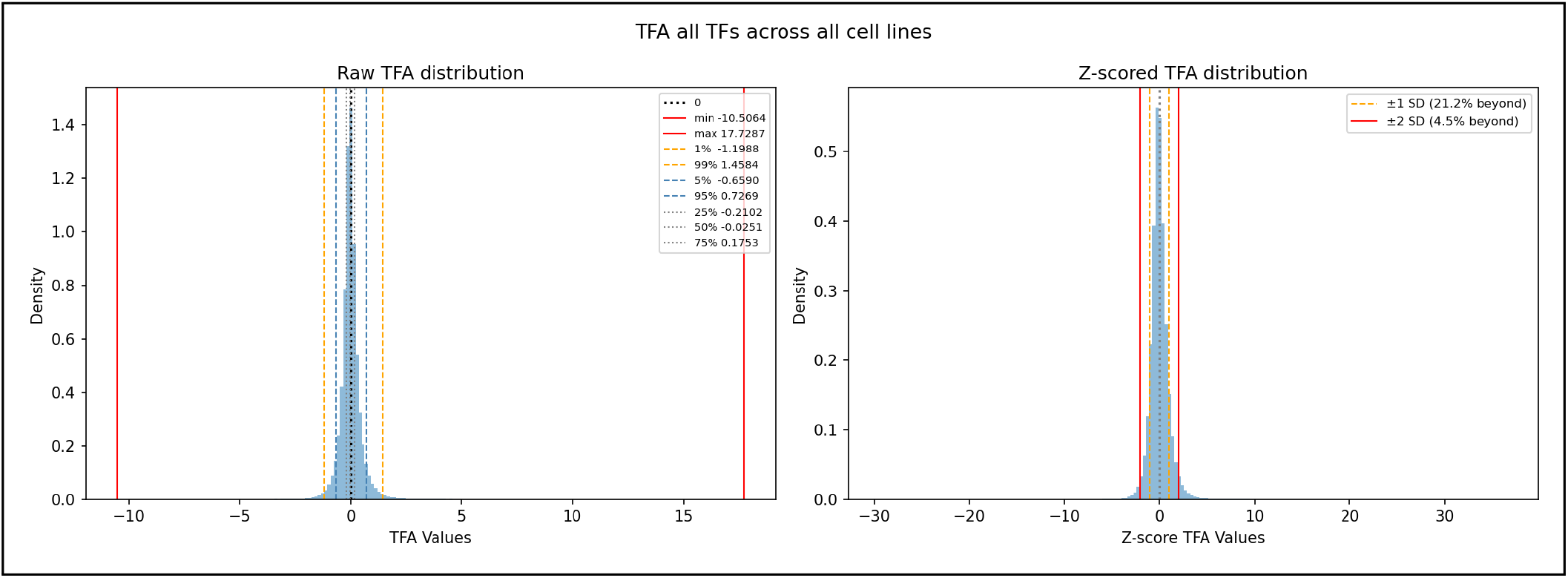
Raw and z-score normalized TFA distributions across all TFs and cell lines. (Left) Raw TFA values range from −10.51 to 17.73; 95% of values fall between −0.66 and 0.73 and 99% between −1.20 and 1.46. (Right) Z-scored TFA values per TF across cell lines; 4.5% of values exceed ±2 standard deviations and are retained as TF input nodes.

Using the defined sources and targets, we construct prizes and actives for each cell line. The magnitudes of the raw TFA values (before z-score normalization) are used as the prize for each retained TF. Somatic mutation prizes are assigned the discrete 0/1/2 coding scheme as scores. Copy number prizes are scored as 0.5. For the actives, we mark all included sources and targets as True.

We combine data that was aligned to different genomes; the CCLE CNA data is aligned to hg19, while the DepMap’s somatic mutation data, TFA prior network, and DepMap’s expression data align to hg38. During the hg19 to hg38 transition, approximately 2% of genes could not be mapped onto the new assembly, and an additional 3% were deleted due to patch consolidations in which two hg19 gene models collapsed into a single hg38 entry.^69^ 95% of gene stable IDs were successfully assigned to models in the new assembly.^69^ We acknowledge that a subset of gene symbols may be mismatched or absent at the intersection of hg19-and hg38-aligned datasets, a limitation inherent to combining data across genome builds.

##### Gold Standard

As a gold standard for evaluation, we use DepMap CRISPR-based gene dependency data,^e^ which summarizes results from genome-wide CRISPR knockout screens across cancer cell lines by testing whether a gene is associated with cell survival or growth. A higher value indicates a greater likelihood that the gene is essential for a given cell line’s survival and proliferation. We select genes for each cell line using a probability threshold of 0.5.

This is not a traditional gold standard, and perfect overlap with reconstructed pathways is not expected. Gene dependency measures whether a gene is essential for survival, whereas pathway reconstruction identifies signaling intermediates connecting sources to targets. A gene may be critical to a signaling pathway without being individually required for cell viability, and conversely, an essential gene may not appear on a reconstructed path between the input sources and targets.

#### 3.2.3 Diseases Collection

This dataset collection captures gene-disease relationships. Complex diseases involve many genes, so reconstruction over disease-associated genes can surface additional relevant genes. Our goal is to assess how well pathway reconstruction methods can recover additional genes associated with those diseases. In this setting, we use target illumination GWAS analytics (TIGA)^71^ confidence scores for disease-associated genes as input node prizes and compare the reconstruction outputs to disease-gene annotations gathered from text mining and knowledge-based evidence.

##### Chosen Diseases

We first identified all disease-gene scores from TIGA and all available disease-gene associations from text mining and curated knowledge channels from the DISEASES 2.0 database.^72^ A disease was included in this collection if it had at least 10 gold standard genes in the DISEASES database and at least one input node prize from TIGA. This yielded 124 disease-specific datasets (Supplementary File 2).

##### Input Nodes

The input nodes for this dataset are GWAS-based disease-gene associations from TIGA.^71^ TIGA calculates confidence scores for gene-trait associations across GWAS, combining citation-based and single nucleotide polymorphism (SNP)-based measures into a mean rank score.^71^ We use only the experimentally-based SNP-based component for the input node prizes. This collection will be used only by algorithms that take prize nodes or active nodes as input because there is no way to designate sources and targets. The TIGA input nodes and STRING text-mining channels share the same underlying node scoring pipeline but score different entity pairs: gene-disease relationships for the input nodes^71^ versus protein-protein relationships for STRING.^43^

##### Gold Standard

The gold standard nodes are derived from the non-experimental sources of evidence from the DISEASES 2.0 database.^72^ The DISEASES database integrates multiple evidence channels (text mining, knowledge, and experiments) to curate gene-disease associations. We construct the gold standard from gene-disease associations that score a four or five on a five-point scale in either the text mining or knowledge channels of DISEASES, excluding the experimental channel because it uses TIGA scores. There is still some overlap between the TIGA input nodes and the DISEASES gold standard nodes with an average of 4 and maximum of 28 nodes across the diseases (Supplementary File 2).

#### 3.2.4 EGFR Phosphoproteomics Collection

This dataset collection contains one omics dataset that captures signaling responses to epidermal growth factor (EGF) stimulation.^73^ Epidermal growth factor receptor (EGFR) is a protein that regulates signaling pathways to control cellular proliferation.^74^ It binds the ligand EGF and triggers several downstream signal transduction cascades.^74^ In this dataset, these events were captured experimentally by stimulating cells with EGF and tracking protein phosphorylation over time using phosphoproteomics.^73^ Phosphoproteomics measures the abundance of phosphorylated peptides using mass spectrometry. Phosphorylation changes reflect shifts in activity and capture molecular responses to stimulation.^75^

##### Input Nodes

The data and processing come from the original phosphoproteomic analysis.^73^ The omics data consists of peptide phosphorylation measurements following EGF stimulation across time points from 0 to 128 minutes. Each peptide was assigned a score reflecting the strength of phosphorylation response to stimulation (scored as −log_10_(*p-value*)), with larger scores indicating smaller p-values. Scores were aggregated by taking the maximum across time points and peptides mapped to a single protein.^73^ We used these protein scores as the input prize, and we mark all proteins with a prize as active. EGF is the experimental stimulus and initiating protein in EGFR signaling, so it is designated as the source node and assigned a fixed prize of 10, greater than all other nodes, to anchor reconstruction at the site of stimulation. EGF is not observed in the phosphoproteomics data, so we explicitly add it as a source, a prize, and an active node. The phosphorylated proteins are assigned as targets, representing downstream pathway components.

##### Gold Standard

The gold standard for this dataset consists proteins in EGFR pathway representations from eight different pathway databases, ^5, 6, 76–80^ which were collected in the original study.^73^ Using the nodes as a gold standard evaluates whether a pathway reconstruction algorithm recovers proteins known to participate in EGFR signaling. However, the pathways give incomplete coverage of the EGF response,^73^ and we do not have an edge-level gold standard.

### 3.3 Evaluation Metrics

We will evaluate the pathway reconstruction algorithms across several categories of metrics. These metrics quantify how well the algorithms predict the gold standards for the datasets above, characterize the graph properties of the reconstructed pathways, compare the reconstructed pathways across algorithms and parameter combinations, and measure the computational performance of the algorithms.

#### 3.3.1 Pathway Statistics

These pathway statistics characterize graph topological properties of the reconstructed pathways, treating those pathways as undirected graphs. Pathway statistics are calculated using NetworkX,^81^ which does not support mixed graphs. Treating all outputs as undirected is an approximation, but it lets us compare across algorithms in a consistent manner.

##### Number of graph components

We report the number of nodes, edges, and connected components in each reconstructed pathway.

##### Density

Graph density measures how tightly connected a network is, measuring how many edges are present relative to the maximum possible edges. For a pathway with *n* nodes and *m* edges, density is 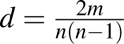 *d* ranges from 0 (no edges) to 1 (complete graph).^82^

##### Node degree

The degree of a node is the number of edges connected to it.^83^ For each reconstructed pathway, we compute the maximum and median degree across all nodes.

##### Diameter

The diameter of a graph is the longest path between any two nodes.^84^ We report the maximum diameter across all connected components in a reconstructed pathway. Singleton components have diameter 0.

##### Average path length

The average path length *a* is the mean shortest path distance over all pairs of distinct nodes, *a* = 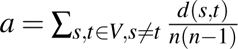, where *d*(*s, t*) is the shortest path between nodes *s* and *t* and *n* is the number of nodes.^85^ We compute this per connected component (excluding singletons) and report the mean across all components.

##### Number of input nodes

For each reconstructed pathway, we report how many nodes were designated as sources, targets, prizes, or dummy nodes in the corresponding input node data.

#### 3.3.2 Gold Standard Evaluation Metrics

A true positive (TP) is a node or edge in the reconstructed pathway that is also in the gold standard. A true negative (TN) is a node or edge absent from the reconstructed pathway that is also absent from the gold standard. A false positive (FP) is a node or edge in the reconstructed pathway that is not in the gold standard. A false negative (FN) is a node or edge absent from the reconstructed pathway that is in the gold standard.^86^

##### Precision

Precision is the proportion of the nodes or edges in the reconstructed pathway that appear in the gold standard, 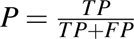.^86^

##### Recall

Recall is the proportion of gold standard nodes or edges recovered in the reconstructed pathway, 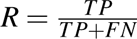.^86^

##### Precision-Recall Curves

A precision-recall curve plots precision against recall across decision thresholds. We generate curves using node and edge frequency scores from the ensembled pathways as thresholds.^86^ Each curve is summarized by its Average Precision, the mean of the precisions at each threshold weighted by the increase in recall, AP = ∑*_n_*(*R_n_* − *R_n_*_−1_) *P_n_,* where *P_n_* and *R_n_* are the precision and recall at the *n*-th threshold.^87^

##### Node-Based Baselines

The first baseline is all nodes in the interactome, which represents the precision of a random predictor. The second baseline is all input nodes, which assesses the overlap between the input nodes and the gold standard nodes. Nodes provided as pathway reconstruction input are not guaranteed to be in the reconstructed pathway graphs, but they are easier to recover. The third baseline is the input nodes plus their first neighbors.^19^

##### Edge-Based Baselines

The first baseline is all edges in the interactome, which represents the precision of a random predictor. The second baseline is all interactome edges induced by the input nodes. That is edges with both endpoints in the input-node set. It measures how much of the gold standard is recoverable directly from the inputs. The third baseline is the set of first-hop edges from the input nodes, meaning every edge with at least one endpoint in the input-node set, respecting directionality so that an edge counts only if it is traversable outward from an input node.^19^

#### 3.3.3 Output Similarity Methods

The following methods operate on a binary edge-by-pathway matrix. Each matrix is dataset-specific. Each row represents an edge in the union of all reconstructed pathways, and each column represents a reconstructed pathway. Each matrix entry indicates whether a given edge appears in a given pathway (1 if present, 0 if absent).

##### Principal Component Analysis (PCA)

PCA is an unsupervised dimensionality reduction technique that transforms high-dimensional data into a lower-dimensional space by projecting it onto orthogonal, linear axes that maximize variance in the data.^88^ We apply PCA to the binary edge-by-pathway matrix and project the data onto the top two principal components (axes), visualizing them in a scatterplot where each point represents a reconstructed pathway. We mean center but do not scale the binary matrix using scikit-learn.^89, 90^

##### Hierarchical Agglomerative Clustering

Hierarchical agglomerative clustering is an unsupervised machine learning technique that recursively builds a hierarchy of clusters. Starting by treating each datapoint (reconstructed pathway) as a separate cluster, hierarchical agglomerative clustering will merge the closet pairs according to a distance metric (Euclidean, Manhattan, or cosine) until all points are combined into a single cluster and displayed as a dendrogram.^91–94^

##### Jaccard Similarity

The Jaccard similarity coefficient of two sets *A* and *B* is <u>^|*A*∩*B*|^</u>, the size of their intersection divided by the size of their union. We compute it between all pairs of reconstructed pathways using their edge sets and display the results as a heatmap.^95, 96^

##### Pathway Ensembling

The pathway ensemble is the union of all reconstructed pathways for a dataset, where each edge is weighted by its mean occurrence across those pathways. Edge frequencies fall in (0, 1]: an edge in every pathway has frequency 1, and edges in no pathway are excluded rather than assigned 0.

#### 3.3.4 Computational Performance Monitoring and Profiling

##### Peak Memory Consumption

Peak memory consumption is the maximum memory consumption during execution, measured in megabytes. Peak memory determines the hardware requirements for running an algorithm on different dataset sizes.

##### CPU time

CPU time is the processor time consumed during execution, measured in microseconds. Unlike runtime, it counts only the intervals when the process is actively executing and excludes I/O waiting and time descheduled by the operating system. We capture it three ways: user CPU time (executing the algorithm’s own code), system CPU time (kernel calls on its behalf), and total CPU time (usually, but not always, the sum of the two).

#### 3.3.5 Visualization

##### Cytoscape

Cytoscape is a software platform for visualizing and analyzing biological networks.^97^ We generate a Cytoscape session file containing visualizations of each reconstructed pathway, allowing users to inspect reconstructed pathways interactively without manually importing results.

### 3.4 Parameter Tuning

The parameter values provided to an algorithm can substantially alter the topological structure of reconstructed pathways and their biological interpretation.^9^ Most algorithms include several parameters with non-intuitive interactions, and values performing well on one dataset may not transfer to another. To select parameter values for each algorithm per dataset, we developed an algorithm-agnostic parameter tuning approach. It consists of a two-stage grid search that uses topological properties of the reconstructed pathways to identify parameter combinations that produce reasonable outputs (Figure 4). This strategy avoids tuning parameters using a gold standard, which risks overfitting to that gold standard, and manual parameter tuning, which introduces subjective bias.

**Figure 4.**
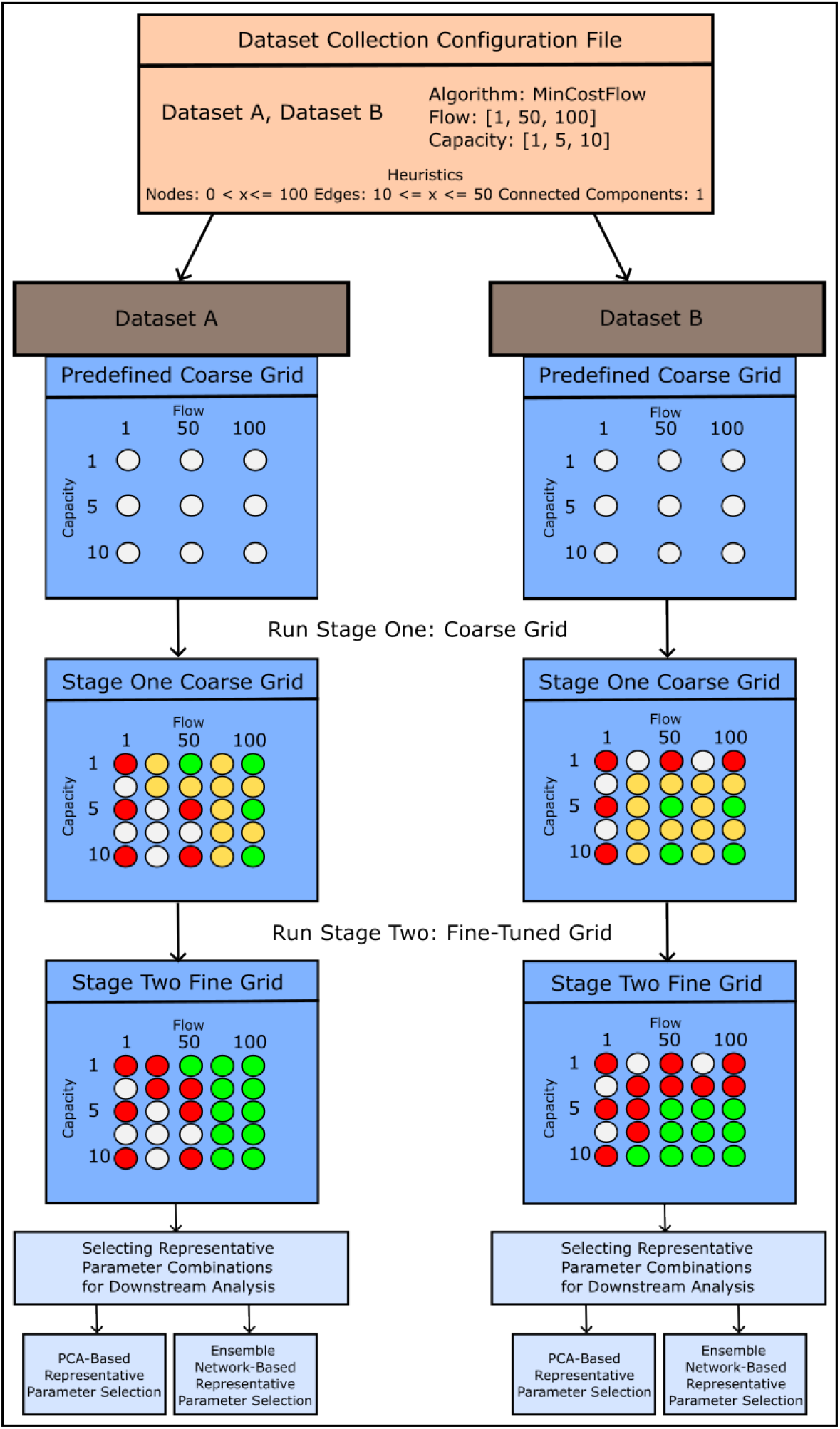
Parameter Tuning: A shared dataset collection configuration defines the algorithms, coarse parameter grid, datasets, and graph heuristics applied across all datasets. Each dataset in a collection is processed independently. During the first stage, the coarse grid is run and evaluated against the heuristics. Parameter combinations whose reconstructed pathways fail the heuristics are discarded (red). During the second stage, candidate midpoint parameter values are generated between each pair of adjacent combinations from the coarse grid, creating the fine-tuned grid. A candidate midpoint is included (yellow) if at least one neighboring coarse grid parameter passes. Midpoints not adjacent to any passing the coarse grid result are ignored (gray). All combinations that pass both stages (green) are then fed into two methods for selecting representative parameter combinations for downstream analysis.

#### 3.4.1 Two-Stage Grid Search

The following strategy (Figure 4) identifies a parameter grid for an algorithm on a specific dataset.

##### Coarse Grid Search

During the first stage, a coarse grid (Supplementary Table 6) of parameter combinations will be run for each algorithm on each dataset within a dataset collection. For the coarse grid, we predefine an initial parameter space for each algorithm that spans a wide range of values with large step sizes and broad lower and upper bounds.

The coarse grid is intended to span a range wide enough to reveal how much an algorithm’s output varies across parameter settings, while remaining small enough to run on all datasets at hand. No general procedure exists for determining ranges for most parameters across algorithms, so ranges were set per algorithm based on the considerations described in Supplementary Section 3.

Each dataset collection has its own configuration file that defines the initial coarse grid and graph heuristics that are used to evaluate reconstructed pathways. These same heuristics are applied across all datasets within a collection rather than being tailored to individual datasets. Each heuristic pairs a graph topological statistic from Section 3.3.1 with a passing range. A parameter combination passes a heuristic the reconstructed pathway from that algorithm with that parameter combination has a graph topological statistic value within the permissible range. A combination fails the value is outside the range or when the algorithm does not return a valid pathway in the time limit. The heuristics encode expectations about plausible reconstructed pathway structure as explicit, reproducible rules. We will choose the statistics and their bounds through a survey of SPRAS team members (Supplementary Section 4).

After running the full coarse grid for each algorithm on each dataset collection, we will evaluate every parameter combination against these heuristics. For each dataset, combinations whose reconstructed pathways pass all the heuristics are kept; all others fail and are not used. Since the underlying biological data vary across datasets within a collection, the set of passing combinations differs per dataset. These per-dataset passing combinations serve as the starting point for the second stage.

##### Fine-Tuned Grid

The passing parameter combinations from the coarse grid will define the start of the fine-tuned grid tailored to each algorithm and dataset. The grid is refined by evaluating the neighbors of passing combinations, where a neighbor is the midpoint between two adjacent coarse-grid values of a single parameter. We treat parameters as independent, varying one at a time to explore the local parameter space more densely.

For each parameter, we define a candidate value at the midpoint between each pair of adjacent coarse-grid values. A candidate midpoint enters the fine-tuned grid when at least one of its two neighboring coarse-grid combinations passed; a candidate between two failing combinations is not evaluated. For example, if the coarse-grid combinations *b* = 5, *d* = 10, *w* = 2 and *b* = 7, *d* = 12, *w* = 4 both pass, we also evaluate the midpoints *b* = 6, *d* = 11, and *w* = 3. If *b* = 5 passes and *b* = 7 fails, the midpoint *b* = 6 is still evaluated. This centers a finer grid on the parameter regions that remain plausible after the coarse grid (Figure 4).

We will apply the same graph heuristics defined for each dataset collection to these neighboring combinations. All combinations that pass, from both the coarse grid and the fine-tuned grid, will form the final parameter grid for each dataset and algorithm.

#### 3.4.2 Selecting Representative Parameter Combinations for Downstream Analysis

The two-stage grid search will yield a set of passing parameter combinations for each dataset and algorithm. From these sets, we will select representative parameter combinations to do downstream analysis and comparisons (Figure 4). SPRAS supports two different approaches to identify representative parameter combinations that enable fair comparisons across algorithms. Each approach makes different assumptions about what constitutes a representative parameter combination and evaluates the resulting outputs differently.

##### PCA-Based Representative Parameter Selection

The PCA approach will select a single representative parameter combination for each pathway reconstruction algorithm on each dataset. For every parameter combination that passes the heuristic, the corresponding reconstructed pathways are projected onto algorithm-specific PCA spaces.

Within this space, we estimate the density of outputs using kernel density estimation and identify the location of the highest density peak. The output closest to this peak by Euclidean distance is selected as the most representative. This approach conceptually seeks a median-like pathway that is most characteristic of the algorithm’s overall behavior on that dataset. Algorithms differ in how many parameter combinations they support, giving some more chances than others to produce a high-performing output. Selecting a single representative per algorithm normalizes this difference. We will then evaluate the selected output against the dataset’s gold standard to compute precision and recall.

##### Ensemble Network-Based Representative Parameter Selection

The ensemble network-based approach will aggregate outputs across all parameter combinations for each pathway reconstruction algorithm. All outputs that pass the heuristic filters will be combined into an algorithm-specific ensemble network. These ensemble networks summarize the topological structures and edges that consistently appear across parameter combinations, which is a common analysis strategy for some algorithms.^98, 99^ For each algorithm-specific ensemble network, we will generate a precision-recall curve by thresholding the edge frequencies.

### 3.5 SPRAS

SPRAS is a software framework for pathway reconstruction that addresses the challenge of running and comparing different algorithms on the same dataset. Users specify datasets, algorithms, parameters, and evaluations in a single configuration file, and SPRAS automatically executes all combinations of the specified inputs. The framework converts SPRAS-formatted inputs into algorithm-specific formats, runs each algorithm in a container, and standardizes algorithm-specific outputs into the SPRAS format. SPRAS leverages Snakemake^11^ for workflow orchestration and uses containers to isolate algorithm dependencies to execute the algorithms across different computing environments. It supports the 14 algorithms, all evaluation metrics and visualizations, and parameter tuning described in Sections 3.1, 3.3, and 3.4.

#### 3.5.1 Data Formats and Conversions

SPRAS requires all input nodes, interactomes, and gold standard datasets to follow a standardized format. This standardization allows SPRAS to convert a single user-provided dataset into algorithm-specific formats. Adding a new dataset requires formatting data once instead of once per algorithm. Pathway reconstruction algorithms also produce outputs in algorithm-specific formats. SPRAS automatically converts these raw outputs into a standardized format to support common post-processing, such as evaluation. Examples are provided in Supplementary Tables 7-13.

SPRAS supports the following node conversions:

- Prizes to actives: if a user provides only prizes, SPRAS defines a binary active set by marking those prize nodes as True.
- Actives to prizes: if a user provides only actives, SPRAS converts them to prizes by assigning each active node a uniform score (e.g., 1.0) to indicate it is a node of interest.
- Source-target to prizes and actives: if a user provides only sources and targets, SPRAS maps sources and targets into prizes by assigning them a uniform score (e.g., 1.0) and into actives by marking them as True.
- SPRAS cannot convert prizes or actives to sources and targets. Prizes and actives alone do not implicitly define which nodes should be treated as upstream starting points versus downstream endpoints.

SPRAS supports the following directionality-aware edge conversions:

- Conversion to undirected edges: Directed edges are converted to undirected by marking them as undirected, ordering edge endpoint identifiers in a canonical order (numerically, then alphabetically), and then removing any duplicated reverse-direction edges.
- Conversion to directed edges: Undirected edges are converted to directed by replacing each edge with two bidirected edges and removing any resulting redundant edges.
- If an algorithm outputs only nodes, SPRAS constructs the induced subgraph over those nodes from the input interactome.

#### 3.5.2 Algorithm Wrapping in SPRAS

One of the primary obstacles to creating a systematic benchmark has been the wide range of programming languages, software dependencies, operating system restrictions, and data formats used across pathway reconstruction algorithms. To address this, each algorithm within SPRAS is wrapped in its own open container image-compliant image. The image contains a software environment packaged with the algorithm’s dependencies and its implementation. This containerization eliminates algorithm dependency conflicts and enables execution across various computing environments. These images can be run with either Docker^100^ or Apptainer.^101^

Algorithm wrapping also includes handling each algorithm’s input and output file formats. SPRAS has wrapper functions to convert input datasets into the format required by each algorithm, including appropriate node types and edge directionality. After an algorithm generates a reconstructed pathway in its raw output format, a wrapper function converts that output into SPRAS’s standardized format.

#### 3.5.3 SPRAS Workflow

SPRAS takes in a user-specified YAML configuration file that defines algorithms, datasets, gold standards, analysis settings, and other settings to control the platform-specific information (Figure 5). SPRAS validates the configuration against a Pydantic schema before running the algorithms, catching many configuration errors before the workflow is executed.^102^

**Figure 5.**
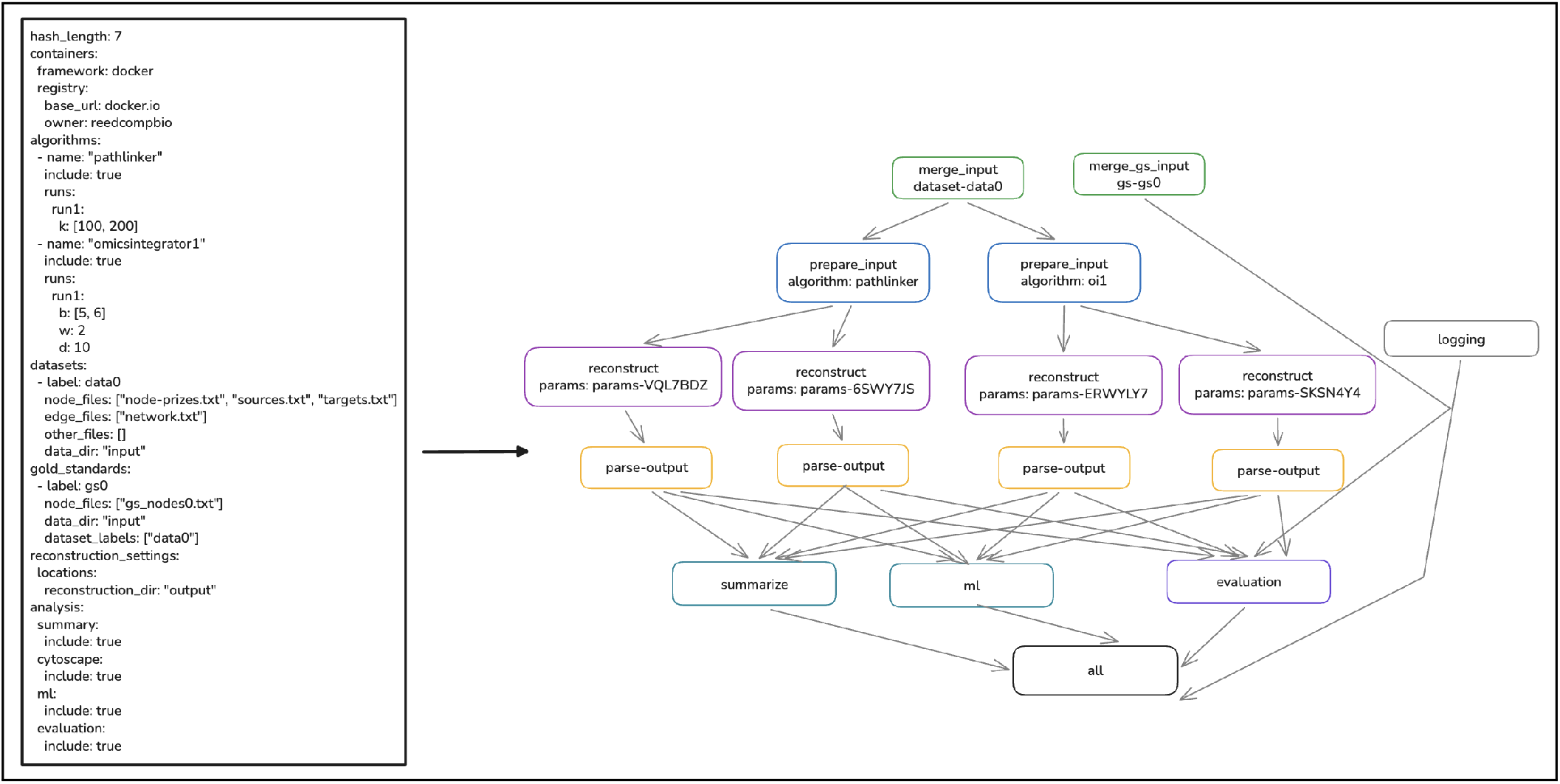
**Left:** Example SPRAS configuration file specifying two algorithms (PathLinker and Omics Integrator 1), each with two parameter combinations, and one dataset with an associated gold standard. **Right:** Snakemake directed acyclic graph generated from the SPRAS configuration file. Each node represents an instantiation of a rule. An edge indicate dependencies between rules. The workflow merges dataset and gold standard inputs, prepares algorithm-specific inputs for PathLinker and Omics Integrator 1, runs reconstruction for each parameter combination (two per algorithm, identified by a unique parameter hash), logs what parameters are associated with each parameter hash, logs what input files are associated with each dataset, parses the output into the common format, and runs downstream analyses (summarization, machine learning, and evaluation).

The central part of the configuration is algorithms and datasets. Users specify which algorithms to run, parameter combinations to apply to each algorithm, and datasets to use. Any dataset specified in the configuration file will have every algorithm and parameter combination run on it. SPRAS executes the tasks set in a configuration file with a Snakemake workflow. Snakemake is a workflow management system that defines computational pipelines as directed acyclic graphs of tasks.^11^ The system manages task dependencies, enabling parallel execution of tasks when those tasks’ prerequisites are met. It also scales execution according to the configuration and tracks completed outputs. When a user updates the configuration or resumes a failed execution, Snakemake recomputes only affected tasks while preserving previously completed work.

Using Snakemake, SPRAS runs all combinations of datasets, algorithm, and parameters provided in a configuration file (Figure 5). For each algorithm specified in the configuration, Snakemake creates tasks to prepare algorithm-specific input data. Then, for each algorithm-parameter-dataset combination, Snakemake creates a task to execute the algorithm in a container. Finally, for each raw output produced by the algorithms, Snakemake creates tasks to transform the outputs into the standardized SPRAS format. If downstream analysis or evaluation tasks (such as any of the metrics mentioned in Section 3.3) are specified in the configuration file, Snakemake creates additional tasks to perform these analyses on the standardized SPRAS outputs.

#### 3.5.4 Configuring Parameter Tuning

Parameter tuning will be integrated directly into SPRAS. Users can specify parameter ranges in the configuration file, and SPRAS will handle the two-stage tuning procedure and downstream analysis described in Section 3.4.

### 3.6 Pathway Reconstruction Assessments

We will assess pathway reconstruction algorithms through three types of evaluations. Pathway recovery performance assesses how well each algorithm recovers gold standard information. Algorithm similarity assesses how similar algorithms are in their outputs. Computational performance assesses the computational feasibility of using these algorithms. Together, these evaluations will highlight different types of strengths and weaknesses across different biological use cases.

#### 3.6.1 Pathway Recovery Performance

Pathway recovery performance (Figure 6) will examine how well each pathway reconstruction algorithm performs the reconstruction task. Specifically, we will measure how well an algorithm recovers gold standard nodes or edges.

**Figure 6.**
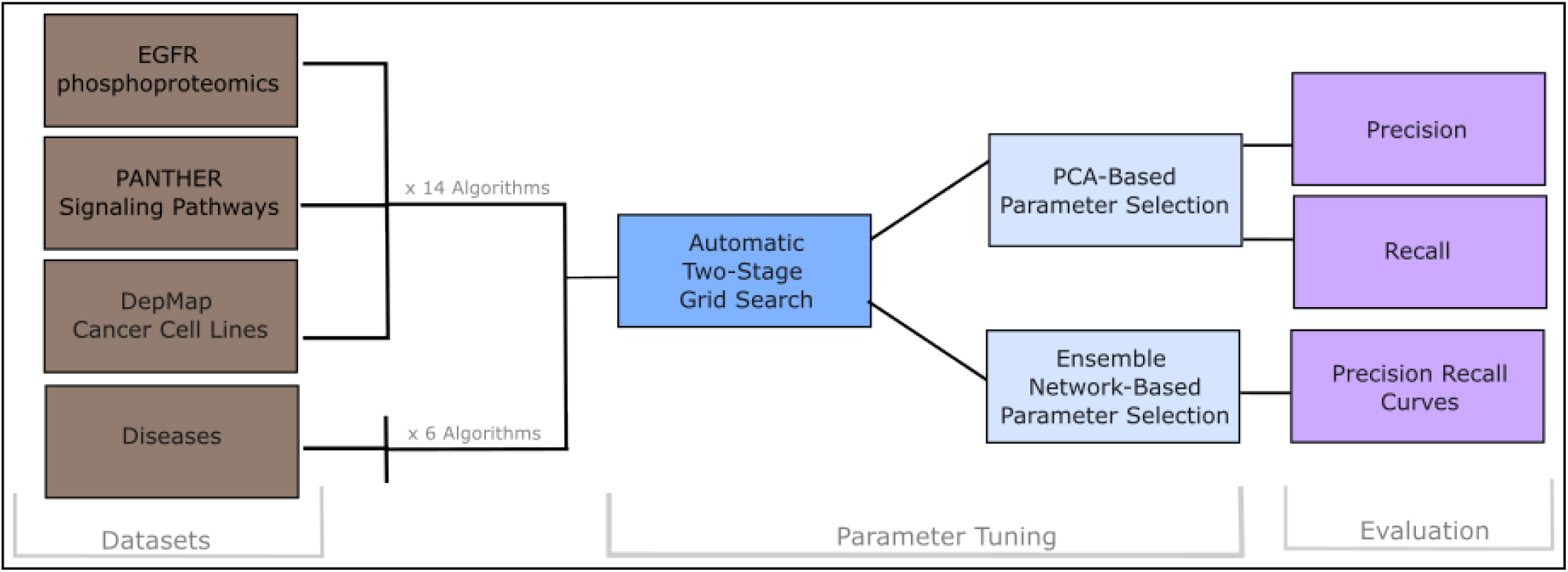
Pipeline for pathway recovery performance assessment. Each dataset collection is analyzed by all wrapped SPRAS algorithms (×14 algorithms for EGFR phosphoproteomics, PANTHER signaling pathways, and DepMap cancer cell lines; ×6 algorithms for DISEASES). For each dataset-algorithm combination, the two-stage grid search constructs a dataset-specific parameter grid. The reconstructed pathways from each grid are then processed by two methods for selecting representative parameter combinations, each running a different downstream analysis against the dataset-specific gold standard.

##### Evaluation Plan

We will use the EGFR dataset collection, the PANTHER pathways collection with the thresholded interactome, the DISEASES collection, and the DepMap cancer cell line collection (Section 3.2). All algorithms (Section 3.1) will be applied to every collection except DISEASES, which is compatible only with algorithms that use prizes or active nodes (Section 3.2.3). For each dataset, we will run the two-stage grid search procedure to construct dataset-specific parameter grids (Section 3.4.1). These grids will then be processed per algorithm using two strategies (PCA-based and ensemble network-based) to identify representative reconstructed pathways for downstream analysis (Section 3.4.2).

To measure pathway recovery performance, we will use precision, recall, and precision-recall curves (Section 3.3.2). Some algorithms optimize for sparse, high-precision pathways while others favor broader interactome exploration; precision and recall quantify both behaviors. We will compute each metric for every algorithm run on each dataset. The strategy used to identify the representative reconstructed pathways determines which metric applies: PCA-based uses precision and recall, while ensemble network-based uses precision-recall curves (Section 3.4.2).

#### 3.6.2 Algorithm Similarity

Algorithm similarity (Figure 7) will examine how similar the outputs of different pathway reconstruction algorithms are on the same dataset. Patterns of similarity could reveal natural groupings of algorithms that behave alike, providing a data-driven alternative to methodological classifications. Evaluating similarity across biological contexts also could reveal whether an algorithm produces variable reconstructed pathways across parameter choices or consistently converges to similar structured pathways.

**Figure 7.**
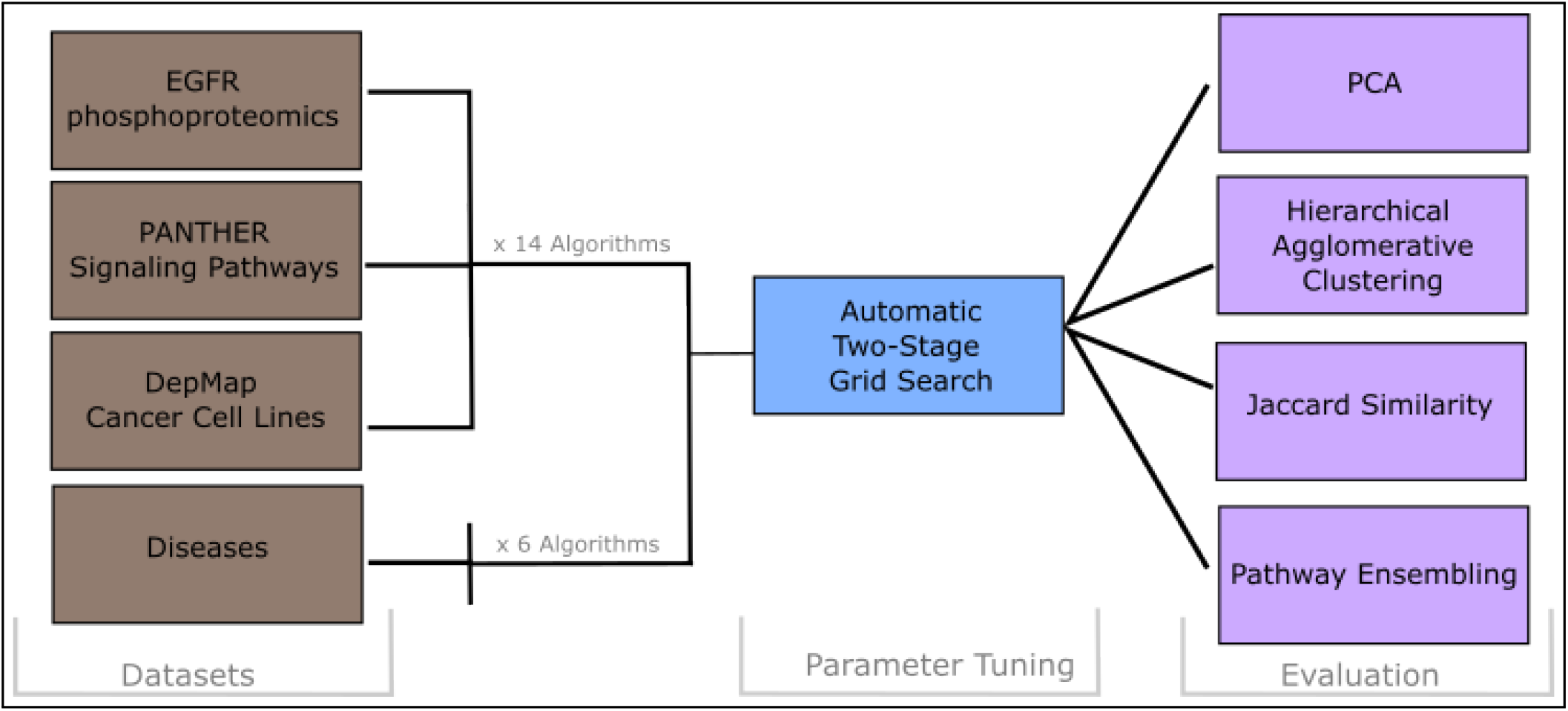
Pipeline for algorithm similarity assessment. Each dataset collection is analyzed by all wrapped SPRAS algorithms (×14 algorithms for EGFR phosphoproteomics, PANTHER signaling pathways, and DepMap cancer cell lines; ×6 algorithms for DISEASES). For each dataset-algorithm combination, the two-stage grid search is run to construct a fine-tuned parameter grid. The resulting reconstructed pathways are then analyzed using four methods to analyze similarity.

##### Evaluation Plan

We will use the EGFR dataset collection, the PANTHER pathways collection with the thresholded interactome, the DISEASES collection, and the DepMap cancer cell line collection (each described in Section 3.2). All algorithms (Section 3.1) will be applied to every collection except DISEASES, which is compatible only with algorithms that use prizes or active nodes (Section 3.2.3). For each dataset, we will run the automatic two-stage grid search procedure (Section 3.4.1) to construct parameter grids per dataset. To assess similarity among algorithms based on their reconstructed pathways, we will apply PCA, hierarchical agglomerative clustering, Jaccard similarity, and pathway ensembling across all algorithms together (Section 3.3.3).

#### 3.6.3 Computational Performance

Pathway reconstruction algorithms optimize objectives over large interactomes and omics datasets, making computational performance a practical concern. By characterizing how algorithms respond to increasing data complexity (interactome size and input nodes varying) and parameter settings, we will provide guidance for selecting algorithms based on available computational resources and dataset characteristics (Figure 8).

**Figure 8.**
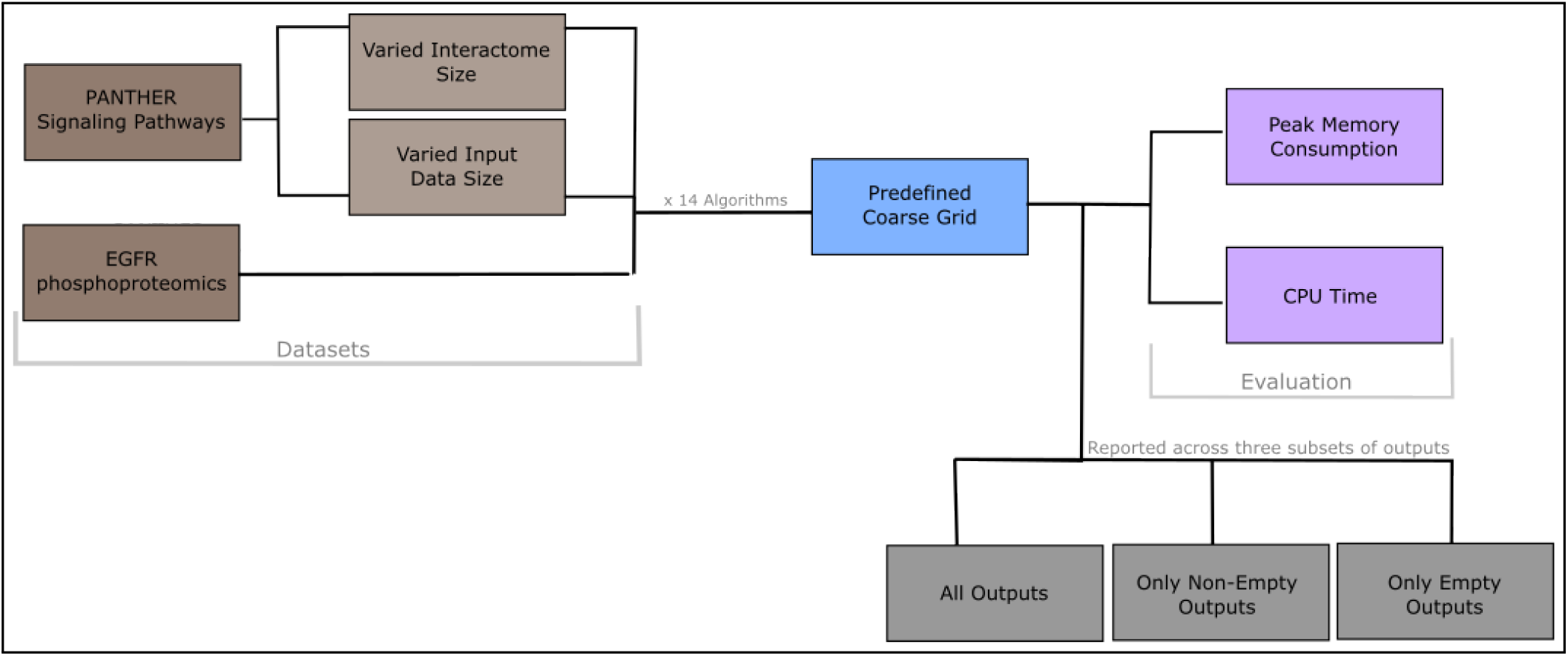
Pipeline for computational performance assessment on PANTHER pathways and EGFR phosphoproteomics. All 14 wrapped SPRAS algorithms are applied to each dataset using only the predefined coarse parameter grid (no heuristic-based parameter tuning) (Supplementary Table 6). Three computational metrics are recorded per run: peak memory consumption and CPU time. Each metric is reported as the mean and standard deviation across three output subsets (all outputs, only non-empty outputs, and only empty outputs) to characterize how algorithm behavior shapes computational cost.

##### Evaluation Plan

For this evaluation, we use two dataset collections: PANTHER (Section 3.2.1) with the downsampled interactomes and EGFR (Section 3.2.4). The PANTHER collection is particularly well-suited for computational performance assessment because we have multiple versions that control the size of the input nodes and input background interactome. The EGFR collection will serve as a biological case study, testing whether the computational performance patterns from the PANTHER evaluation transfer to an experimental omics dataset.

From the 22 PANTHER pathways (Supplementary Table 2), we selected 11, chosen by their source/target ratios and grouped into three categories: balanced, source-focused, and target-focused. Balanced pathways have a source-to-target ratio *r* within 0.15 of 1.0 (0.85 ≤ *r* ≤ 1.15), source-focused pathways have a source-to-target ratio *r* ≥ 3.4, and target-focused pathways have a target-to-source ratio of at least 3.4. The complete list is provided in Supplementary Table 14. We restrict to these 11 because they fall cleanlyn into these three categories, letting us assess algorithm performance across distinct pathway structures. Each of the 11 pathways is paired with the five downsampled STRING physical interactomes constructed at 10, 30, 50, 70, and 100 percent (Section 3.2.1). We will also use the four merged pathway variants (Section 3.2.1), each paired with the 100% downsampled STRING physical interactome.

All algorithms will be applied to every selected dataset using the coarse parameter grid defined for each algorithm (Supplementary Table 6), which provides a finite set of parameter configurations that affect memory consumption and CPU time. We will only run the predefined coarse grid; we will not apply the stage-one coarse grid heuristics of the two-stage grid search workflow. The coarse grid gives a sufficient characterization of how these parameters affect performance.

We will run two experiments to isolate the effect of each input data component on computational performance. The first varies the input nodes: we will run all 11 PANTHER pathways and 4 merged pathway variants on the full STRING interactome (100%), holding the network fixed. The second varies the interactome size: we will run the 11 chosen PANTHER pathways across all 5 downsampled interactomes, holding the input nodes fixed.

For each experiment, we will report peak memory consumption and CPU time (Section 3.3.4). For each algorithm, we will visualize the distribution of each performance metric and compute its mean and standard deviation across all interactome-input node combinations and parameter configurations. We will report these statistics separately for all parameter combinations, combinations that produce non-empty outputs, and combinations that produce empty outputs. This decomposition will reveal how parameters and input data affect computational performance across different algorithm behaviors.

All algorithms will be executed on a single CPU core. For algorithms that support parallel processing, this will constrain execution and may underestimate their performance under multi-core usage, but it will ensure a comparable baseline across all algorithms. Each run will also be subject to a maximum CPU time of three days and a memory limit of 64 GB.

### 3.7 Running at Scale

The full benchmark comprises 822 datasets that will be evaluated across 14 pathway reconstruction algorithms. In the worst case, running both the coarse and fine parameter grids, the search spans 4,205 combinations. The DISEASES collection uses a reduced set of 3,952 combinations as it will run only the 6 algorithms that use prizes or active nodes. Of the 822 datasets, the 124 in the DISEASES collection will use the 3,952-combination set, and the remaining 698 will use the full 4,205-combination set, for a worst-case total of 3,425,138 algorithm runs. Accounting for the additional workflow steps required to support these reconstructions (data preparation, parameter and dataset logging, output parsing), the total job count could reach 6,867,464. Further details are provided in Supplementary Table 15, Supplementary Table 16, and Supplementary File 3.

Running this benchmark at this scale will require distributed computation. SPRAS executes through Snakemake, which manages job parallelization and task dependencies. The benchmark will run on the Open Science Pool (OSPool),^103–106^ a national distributed high-throughput computing resource, through the Center for High Throughput Computing (CHTC).^107^ Snakemake dispatches jobs to OSPool through an HTCondor executor plugin, which submits and manages each job on HTCondor. Each algorithm will execute inside an Apptainer^101^ container, built from a corresponding Docker^100^ image, to ensure consistent software environments across heterogeneous OSPool resources. Every Snakemake job will run independently across available OSPool resources, requesting one CPU core, 8 GB of memory, and 16 GB of disk. If a job fails from insufficient memory, it will be retried with its memory request raised by 4 GB per attempt, up to 64 GB.

## 4 Pilot Study

Our pilot study evaluated multiple pathway reconstructions algorithms across multiple datasets and was used to finalize the design decisions for our full scale benchmark.^108^ The pilot datasets consisted of five PANTHER^7^ pathways and three versions of the *Homo sapiens* STRING v12 interactome^43^ thresholded at different experimental score levels. The PANTHER pathways were selected to provide variation in topology, source and target availability, and network scale. Seven pathway reconstruction algorithms were run under broad parameter grids: PathLinker, Omics Integrator 1, Omics Integrator 2, MinCostFlow, Maximum Edge Orientation, All Pairs Shortest Paths, and DOMINO.

The pilot study confirmed that SPRAS could operate at scale, successfully coordinating multiple algorithms across diverse datasets and parameter combinations. However, the pilot exposed workflow and computational bottlenecks that will be addressed in our full-scale benchmark. The pilot run confirmed that the representative parameter-selection methods described in Section 3.4.2 were appropriate: PCA-based and ensemble network-based selection provided complementary views of pathway recovery performance. However, defining parameter ranges proved more complex than initially anticipated. The reconstructed pathway size and structure depended not only on chosen parameter combinations, but also on underlying dataset characteristics and biological context. This revealed the need for a systematic, dataset-specific method to identify parameter grids separately for each dataset and algorithm, rather than applying a single universal grid for all datasets. This finding directly motivated the development of the automatic two-stage grid search approach we will deploy in the full benchmark (Section 3.4.1).

Consistent with our expectations, the pilot study evaluations showed that no single pathway reconstruction algorithm performed best across all datasets. The evaluation visualizations, particularly the precision-recall curves for ensemble networks, were difficult to compare across interactomes, pathways, and algorithms because each was a separate plot, making it hard to draw conclusions across them. These findings highlighted the need for additional datasets. The pilot used a single biological setting, so we could not tell whether performance patterns were general or specific to that context. In the full benchmark, we therefore added datasets spanning multiple biological settings, enabling more context-dependent interpretation of algorithmic performance. They also motivated redesigning evaluation visualizations for easier interpretation and comparison.

Additionally, we spot-checked parameter behavior on a preliminary version of the EGFR dataset that used a different background interactome.^73^ Examining how reconstructed pathways changed across a range of parameter values confirmed that our coarse grids are reasonable. It helped verify chosen values, refine the grid where preliminary behavior indicated missing coverage or redundancy, and build an understanding of parameters that lacked clear documentation.

We also assessed the computational feasibility of running the full coarse grid (not two-stage parameter tuning) on the OSPool. Using the EGFR dataset described for the full benchmark, we estimated the runtime, memory, and disk requirements of each algorithm, which let us calculate the total number of jobs and estimate how long it would take to run the full benchmark (Supplementary File 3).

## 5 Discussion

### 5.1 Network Algorithm Benchmarking Frameworks

Several related software frameworks have addressed similar challenges as SPRAS. SPRAS draws direct inspiration from BEELINE,^109^ a modular benchmarking framework for gene regulatory network (GRN) inference. BEELINE supports running, comparing, and evaluating many GRN methods within a standardized pipeline. It preprocesses datasets, executes Dockerized implementations of multiple GRN methods, and provides downstream post-processing, visualization, and evaluation tools. SPRAS focuses on pathway reconstruction rather than GRN inference and also makes different design decisions. SPRAS uses Snakemake to manage workflow execution, which enables scalable, resumable pipelines. Additionally, SPRAS will include a parameter-tuning feature that helps identify parameter grids for each dataset and then pick representative parameters for downstream analysis.

A recent disease module identification pipeline shares motivation and several high-level design decisions with SPRAS.^19^ Like SPRAS, it uses containers to manage software dependencies and reduce installation challenges, converts between common and algorithm-specific input and output formats, and orchestrates execution dependencies with a workflow manager (Nextflow^110^ rather than Snakemake). It supports six disease module algorithms, three of which SPRAS also provides (DIAMOnD,^20^ DOMINO,^27^ and Random Walks with Restarts^111^). Whereas that pipeline targets disease module identification alone, SPRAS encapsulates both pathway reconstruction and disease module identification within one framework. Another framework similar to SPRAS is NetworkCommons.^12^ It is a Python package for reconstructing biological pathways by integrating prior knowledge networks (interactomes), omics data, and eight pathway reconstruction algorithms within a modular workflow. Each method in NetworkCommons is exposed through a consistent Python API and must be callable from Python, either implemented directly in the package or accessed through a Python wrapper around a supported library. In contrast, SPRAS containerizes each algorithm using Docker, so it can support tools written in any programming language or with conflicting Python dependencies. The two frameworks support largely different sets of algorithms; the only ones both provide are All Pairs Shortest Paths^25^ and Random Walks with Restarts.^111^ Rather than porting every method from their framework, we included a representative set spanning the methodological categories described in Section 2.1. NetworkCommons also requires users to write Python code to configure an evaluation pipeline. SPRAS instead uses workflow automation, where users specify settings in a configuration file and Snakemake handles pipeline assembly and execution.

### 5.2 Additional Pathway Reconstruction Algorithm Evaluations

There are other relevant benchmarking studies for pathway reconstruction,^3, 13^ disease module identification,^19, 112–116^ and related network inference^117, 118^ that place less emphasis on a software framework. They also treat pathway reconstruction and disease module identification as separate network biology problems.

For pathway reconstruction, a prior benchmark evaluated seven interactomes and four pathway reconstruction algorithms on known pathways, finding that interactome choice substantially affected reconstruction performance.^13^ Another benchmark examined signaling pathway inference from phosphoproteomics data, evaluating three reconstruction methods across multiple kinase-substrate resources and datasets focused on the EGF response. They found that interactomes choice had greater influence on recovered interactions than either the reconstruction method or experimental context.^3^

Several benchmarks have assessed disease module identification. Studies of four to 75 methods, spanning many biological datasets and interactomes, reported that no single method was best and that results varied with interactome and algorithm choice, input signal strength, and evaluation metric.^19, 112, 113, 116^ A separate study compared eight methods on real PPI networks against degree-preserving random networks and found that most produced no more biologically meaningful modules on the real networks, indicating that they mainly exploit node degree rather than the biological information encoded in edges.^115^

In the related task of topology-based pathway analysis, a benchmark compared seven methods and examined how parameters affect the results, reporting wide variability across methods and providing recommendations for selecting a method for a given analysis task.^117^ Another benchmark studied compound mechanism of action based on how targets and pathways could be recovered from gene expression effects. It evaluated four algorithms paired with four prior-knowledge networks across 269 compounds and found that the combination of algorithm and interactome most determined recovery.^118^ A network module evaluation assessed how shallow and deep graph representation learning methods impacted module quality.^114^

### 5.3 Pathway Reconstruction Parameter Tuning

The parameters required for most pathway reconstruction algorithms can produce reconstructed pathways with drastically different topological properties and biological interpretations.^9^ Parameters may assign weights to nodes or edges, define algorithmic tradeoffs, or specify the search space size. Pathway reconstruction parameter tuning faces irregular parameter landscapes, dataset-dependent performance, and high computational costs from running an algorithm under many parameter combinations.

Several parameter tuning strategies for pathway reconstruction resemble ML hyperparameter optimization techniques.^119^ Metric-based approaches use measures such as F1 score, area under the precision-recall curve, precision, and recall evaluated against gold standard data to identify suitable parameters.^13, 38, 109, 120^ Evaluations based on gold standard data are an inappropriate way to select parameters for pathway reconstruction in general. Pathway reconstruction is typically an exploratory analysis used to summarize data and generate hypotheses,^9^ and there is rarely a gold standard available. Cross-validation can be used to identify parameters that best recover held out input data,^9^ but methods perform poorly on this task.^19^ In addition, algorithms vary in how they define the optimization objective, so no single intrinsic metric can serve as a universal tuning target.

Alternative approaches focused on network topology rather than metrics. DEGAS compares the size of the reconstructed pathway to a background distribution of sizes of reconstructed pathways obtained when the interactome node identities are shuffled.^121^ Parameter advising selects settings by identifying reconstructed pathways whose topological features resemble curated biological pathways using graphlets.^9^ While algorithm-agnostic, this approach is limited to undirected graphs and requires substantial computational overhead. Other approaches use graph properties to select parameters for a single dataset,^73, 98, 99, 122^ choosing parameter ranges and graph properties that are suitable for that dataset.

Some pathway reconstruction applications rely on manual, subjective tuning.^9^ Manual tuning introduces the risk of human error and inadvertent bias in the reconstructed pathways and limits the number of parameter combinations that can be assessed.^9^ Manual tuning is further complicated when algorithms’ parameters are not well documented or lack guidance about appropriate ranges. The two-stage grid search approach that we will use in our benchmark represents a compromise. It requires a manual, user-defined coarse grid but then selects reconstructed pathways and fine-tuned gride parameter combinations objectively without using a gold standard.

### 5.4 Limitations and Future Work

#### 5.4.1 PCA on Binary Data

SPRAS applies PCA to a binary data matrix, where each entry indicates whether an edge is present or absent in a reconstructed pathway. Standard PCA assumes normally-distributed data^123^; applying it to binary data violates this assumption. Despite this, PCA on non-Gaussian data is a common practice.^124^ Logistic PCA is designed specifically for dimensionality reduction on binary data, as it models Bernoulli rather than Gaussian distributions.^123, 124^ We will assess logistic PCA as a replacement for PCA based on the similarity of the low-dimensional pathway representations from both methods and the scalability of logistic PCA.

#### 5.4.2 Licensing Issues for Wrapped Algorithms

SPRAS wraps pathway reconstruction algorithms as containers to support execution across diverse computational environments. However, not all pathway reconstruction algorithms can be included in SPRAS due to licensing incompatibilities. Some relevant algorithms are distributed under licenses that prohibit redistribution. As a result, the algorithms supported in SPRAS are constrained by what is permitted under each algorithm’s license. Expanding algorithm coverage in future versions of SPRAS will require either identifying open source alternatives, obtaining suitable licenses from algorithm authors, or reimplementing these methods.

#### 5.4.3 Reimplementation of Pathway Reconstruction Algorithms

No public implementation exists for some algorithms included in SPRAS, requiring reimplementation based on the algorithm descriptions provided in the original publications. In other cases, public code exists but contains bugs or compatibility issues, so we create patches in repository forks or apply them when building the Docker image. As a result, some algorithms in SPRAS are not the exact original implementations but are instead our best approximations. For further details, refer to Supplementary Table 1.

#### 5.4.4 Interactome Choice

This benchmark evaluates pathway reconstruction algorithms using a single interactome. Interactome choice can affect algorithm performance, as differences in edge weight distributions, protein coverage, network density, evidence composition, and directionality conventions influence reconstruction out-comes.^13, 19, 22^ Interactomes are additionally subject to study bias, skewing toward well-studied molecules and interactions due to the tendency of research to concentrate on highly characterized proteins, which can influence which nodes and edges algorithms select independent of biological relevance.^13, 22^ Our benchmark’s scope is on algorithm differences rather than interactome quality. Other work has begun to address this question systematically,^3, 13, 22^ and evaluating how different interactomes affect pathway reconstruction algorithms is an avenue for future work.

#### 5.4.5 Gold Standards

Pathway reconstruction lacks comprehensive gold standards in most cases. Our evaluations mainly use node-level gold standards, except for the PANTHER pathways where the goal is to recover pathway edges from partial pathway nodes. A node-level gold standard evaluates whether the reconstructed pathway identifies a set of biologically relevant molecules, not whether the reconstructed edges or overall pathway structure are correct. There are valid biological reasons why some of the nodes in one of our gold standards may not appear in a pathway and why pathway members may not appear in that gold standard. For instance, the CRISPR-based gene-dependency scores^51, 70^ used as the gold standard for the cancer cell line DepMap datasets can contain false positives or false negatives due to relevant genes that do not impact cell proliferation or death,^125^ synthetic lethality,^125, 126^ limitations of cells in a 2D monolayer as a cancer model,^50, 125, 127^ and off-target effects or other subtle data biases that persist after normalization.^50, 125^

## 6 Timeline

We anticipate a timeline of approximately eight months from approval of the registered report to completion of the final manuscript (Table 1). In the worst case, where every parameter combination in the fine-tuned grid is run on each dataset, we estimate executing the benchmark will require 195 days of wall time across 200 parallel CPUs (Supplementary File 3). The actual execution time will depend on how many OSPool CPUs we can access on average, so it could range from one to six months. Based on our pilot runs with the OSPool, we expect the benchmark to complete closer to one month. However, we allot three months total to leave room for any issues that may occur.

**Table 1.** Timeline.

| Month | Tasks |
| --- | --- |
| 1-3 | Finish code and dataset setup |
| 3-6 | Run the benchmark in the OSPool |
| 6-8 | Interpret results and write updated manuscript |

**Table 2.** Estimated Wall Time.

| # of CPUs | Wall time (days) |
| --- | --- |
| 200 | 195 |
| 500 | 78 |
| 1,000 | 39 |

## 7 Acknowledgments

This material is based upon work supported by the National Science Foundation under awards 2233968 (to A.G.) and 2233969 (to A.R.). S.A.H is supported by National Institutes of Health awards R01GM117339, R01GM144708, and T15LM007359. N.P. is supported by National Institutes of Health award U01HG012039 and the University of Wisconsin–Madison Office of the Vice Chancellor for Research and Graduate Education with funding from the Wisconsin Alumni Research Foundation. This research will be executed through CHTC^107^ using services provided by the OSG Consortium,^103–106^ which is supported by the National Science Foundation awards 2030508 and 2323298.

We thank Margaret Elliott for a bug fix and the initial ensembling code, Sabah Hoq Khondaker for analyzing predicted interactomes, Emma Graham Linck for testing Omics Integrator 2 to inform future SPRAS integration, Thu Ngo for work on the BowTieBuilder implementation, Tobias Rubel and Pramesh Singh for conceptual feedback, Matt Stefely for helping create the overview figure, Carol Sze for extending the network summarization code, Alicia Williams for reviewing and editing the manuscript, Zhiqian Xu for investigating network algorithms for suitability in SPRAS, Sam Yeleti for work on the BowTieBuilder implementation, and Nina Young for work on the All Pairs Shortest Paths implementation.

## 8 Author contributions statement

**Code** N.T., T.F.R., J.H., C.S.M., A.S., N.P., Y.L., S.S., O.F.A., A.B., O.T.J., J.A.H., A.N., M.D., C.L., G.H., A.R., A.G.

**Documentation** N.T., T.F.R., J.H., M.D.

**Data collection and processing** N.T., T.F.R., Y.L., S.S., O.F.A., A.B., A.O., S.A.H., I.J., D.N., G.H.L., A.R., A.G.

**Analysis** N.T.

**Writing** N.T., T.F.R., A.N., I.J., A.R., A.G.

**Figures** N.T.

**Literature Searches** N.T., A.O., A.R., A.G.

**Manuscript Review** N.T., T.F.R., J.H., C.S.M., A.S., N.P., Y.L., S.S., O.F.A., A.B., A.O., O.T.J., J.A.H.,

S.A.H., A.N., I.J., M.D., D.N., C.L., G.H., G.H.L., A.R., A.G.

**Supervision** N.T., O.F.A., A.B., A.R., A.G.

## 9 Data and Code Availability

SPRAS is available under an MIT license.

**SPRAS framework code:** https://github.com/Reed-CompBio/spras, archived at https://doi.org/10.5281/zenodo.13366795.

**Dataset processing code and data:** https://github.com/Reed-CompBio/spras-benchmarking

**SPRAS documentation:** https://spras.readthedocs.io/

## Footnotes

a OmicsSomaticMutationsMatrixDamaging.csv DepMap 2025Q3^51^

b EstimateNCA^63,64^

c OmicsExpressionTPMLogp1HumanAllGenesStranded.csv DepMap 2025Q3^51^

d file_inputs/mammal_file_inputs/hesc/prior.txt.gz^63,66^

e CRISPRGeneDependency.csv DepMap2025Q3^51,70^

