## Supplementary Information for "A Framework for Benchmarking Pathway Reconstruction Algorithms"

### Supplementary Information: A Framework for Benchmarking Pathway Reconstruction Algorithms

#### 1 Algorithm Details

| Algorithm | Algorithm Type | Input Nodes | Input Dir. | Output Dir. | Uses Edge Weights | Parameters | Implementation |
|---|---|---|---|---|---|---|---|
| All Pairs Shortest Paths[1] | Shortest paths | Sources and Targets | Both | Undirected | Yes | N/A | Reimplementation using NetworkX |
| BowTieBuilder[2] | Shortest paths | Sources and Targets | Directed | Directed | Yes | N/A | Reimplementation |
| DIAMOnD[3] | Module detection | Actives | Undirected | Undirected | No | $n$, $\alpha$ | Original code |
| DOMINO[4] | Module detection | Actives | Undirected | Undirected | No | slice threshold, module threshold | Original code |
| Maximum Edge Orientation[5] | Edge orientation optimization | Sources and Targets | Both | Directed | Yes | max path length, local search, rand restarts | Original code |
| MinCostFlow[6–8] | Network flow | Sources and Targets | Both | Both | Yes | flow, capacity | Reimplementation using OR-Tools |
| NetMix2[9] | Network propagation | Prizes | Undirected | Undirected | No | $\delta$, density, num_edges | Original code |
| Omics Integrator 1[10] | Steiner forest | Prizes and Dummy Nodes | Both | Both | Yes | $w$, $b$, $d$, $\mu$, $g$, $r$, noisy_edges, noise, random_terminals, shuffled_prizes, seed, dummy_mode | Original code |
| Omics Integrator 2[11] | Steiner forest | Prizes and Dummy Nodes | Undirected | Undirected | Yes | $w$, $g$, $b$, noise, noisy_edges, random_terminals, dummy_mode | Modified fork of original code |
| PathLinker[12,13] | Shortest paths | Sources and Targets | Directed | Directed | Yes | $k$ | Original code |
| Random Walks with Restarts[14] | Random walk | Prizes | Directed | Directed | Yes | threshold, $\alpha$ | Reimplementation |
| ResponseNet[15] | Network flow | Sources and Targets | Directed | Undirected | Yes | $\gamma$ | Reimplementation |
| Source-Target Random Walks with Restarts[14] | Random walk | Sources and Targets | Directed | Directed | Yes | threshold, $\alpha$ | Reimplementation |
| TieDIE[16] | Network propagation | Sources and Targets | Undirected | Undirected | No | $s$, $a$, $c$, $p$, pagerank, all_paths | Modified fork of original code |

**Supplementary Table 1.** 14 Pathway reconstruction algorithms wrapped in SPRAS.

#### 2 PANTHER pathways

| PANTHER Pathway | # of sources | # of targets | Source-target ratio | Target-source ratio |
|---|---|---|---|---|
| Apoptosis signaling pathway | 8 | 17 | 0.471 | 2.125 |
| B cell activation | 5 | 2 | 2.500 | 0.400 |
| Cadherin signaling pathway | 104 | 3 | 34.667 | 0.029 |
| EGF receptor signaling pathway | 12 | 7 | 1.714 | 0.583 |
| FAS signaling pathway | 2 | 1 | 2.000 | 0.500 |
| Gastrin_CCK2R_240212 | 4 | 20 | 0.200 | 5.000 |
| Hedgehog signaling pathway | 2 | 2 | 1.000 | 1.000 |
| Hypoxia response via HIF activation | 1 | 3 | 0.333 | 3.000 |
| Inflammation mediated by chemokine and cytokine signaling pathway | 31 | 9 | 3.444 | 0.290 |
| Insulin/IGF pathway-mitogen activated protein kinase kinase/MAP kinase cascade | 4 | 2 | 2.000 | 0.500 |
| Interferon-gamma signaling pathway | 2 | 5 | 0.400 | 2.500 |
| Interleukin signaling pathway | 18 | 16 | 1.125 | 0.889 |
| JAK/STAT signaling pathway | 1 | 9 | 0.111 | 9.000 |
| Notch signaling pathway | 13 | 3 | 4.333 | 0.231 |
| p38 MAPK pathway | 1 | 6 | 0.167 | 6.000 |
| PDGF signaling pathway | 2 | 28 | 0.071 | 14.000 |
| PI3 kinase pathway | 2 | 3 | 0.667 | 1.500 |
| T cell activation | 6 | 2 | 3.000 | 0.333 |
| TGF-beta signaling pathway | 12 | 14 | 0.857 | 1.167 |
| Toll receptor signaling pathway | 9 | 3 | 3.000 | 0.333 |
| VEGF signaling pathway | 1 | 2 | 0.500 | 2.000 |
| Wnt signaling pathway | 106 | 15 | 7.067 | 0.142 |

**Supplementary Table 2.** 22 chosen PANTHER signaling pathways.

| Merged Variant Pathways | # of sources | # of targets | Source-target ratio | Target-source ratio |
|---|---|---|---|---|
| Mega pathway | 229 | 98 | 2.337 | 0.428 |
| Source-focused pathway | 84 | 38 | 2.211 | 0.452 |
| Target-focused pathway | 44 | 70 | 0.629 | 1.591 |
| Balanced pathway | 98 | 86 | 1.140 | 0.876 |

**Supplementary Table 3.** Four merged variant pathways. The mega pathway combines all 22 pathways into one. The target-focused pathway merges all pathways with more than 10 targets (excluding the Wnt pathway, which would dominate the source side), and the source-focused pathway merges all with more than 10 sources (excluding Cadherin and Wnt pathways). The balanced pathway is built by greedy expansion.

| PANTHER Pathway | Total edges | | | | |
|---|---|---|---|---|---|
| | $t = 10$ | $t = 30$ | $t = 50$ | $t = 70$ | $t = 100$ |
| Apoptosis signaling pathway | 74,147 | 221,900 | 369,637 | 517,367 | 739,007 |
| B cell activation | 74,017 | 221,772 | 369,512 | 517,253 | 738,898 |
| Cadherin signaling pathway | 76,485 | 224,214 | 371,924 | 519,645 | 741,256 |
| EGF receptor signaling pathway | 75,448 | 223,164 | 370,868 | 518,583 | 740,182 |
| FAS signaling pathway | 73,928 | 221,689 | 369,441 | 517,189 | 738,840 |
| Gastrin_CCK2R_240212 | 74,089 | 221,838 | 369,580 | 517,319 | 738,952 |
| Hedgehog signaling pathway | 73,942 | 221,704 | 369,450 | 517,201 | 738,855 |
| Hypoxia response via HIF activation | 73,904 | 221,665 | 369,414 | 517,167 | 738,821 |
| Inflammation mediated by chemokine and cytokine signaling pathway | 75,863 | 223,548 | 371,207 | 518,894 | 740,446 |
| Insulin/IGF pathway-mitogen activated protein kinase kinase/MAP kinase cascade | 73,963 | 221,722 | 369,471 | 517,221 | 738,867 |
| Interferon-gamma signaling pathway | 73,970 | 221,728 | 369,475 | 517,227 | 738,876 |
| Interleukin signaling pathway | 74,671 | 222,402 | 370,128 | 517,847 | 739,463 |
| JAK/STAT signaling pathway | 73,938 | 221,701 | 369,450 | 517,203 | 738,857 |
| Notch signaling pathway | 73,975 | 221,730 | 369,477 | 517,227 | 738,871 |
| p38 MAPK pathway | 73,932 | 221,690 | 369,438 | 517,186 | 738,834 |
| PDGF signaling pathway | 74,591 | 222,328 | 370,054 | 517,783 | 739,411 |
| PI3 kinase pathway | 74,056 | 221,811 | 369,555 | 517,301 | 738,941 |
| T cell activation | 74,118 | 221,864 | 369,603 | 517,320 | 738,948 |
| TGF-beta signaling pathway | 74,940 | 222,666 | 370,396 | 518,105 | 739,716 |
| Toll receptor signaling pathway | 73,955 | 221,712 | 369,459 | 517,203 | 738,846 |
| VEGF signaling pathway | 74,048 | 221,803 | 369,546 | 517,293 | 738,944 |
| Wnt signaling pathway | 76,480 | 224,156 | 371,796 | 519,445 | 740,988 |

**Supplementary Table 4.** Total number of edges in the downsampled physical interactomes for the 22 PANTHER signaling pathways at downsampling thresholds $t \in \{10, 30, 50, 70, 100\}$, where $t$ is the percentage of the interactome retained. The total number of edges is the pathway plus interactome network at each threshold.

| PANTHER Pathway | Pathway-interactome overlap | | | | |
|---|---|---|---|---|---|
| | $t = 10$ | $t = 30$ | $t = 50$ | $t = 70$ | $t = 100$ |
| Apoptosis signaling pathway | 0.103 | 0.249 | 0.324 | 0.442 | 0.617 |
| B cell activation | 0.132 | 0.289 | 0.353 | 0.542 | 0.753 |
| Cadherin signaling pathway | 0.021 | 0.057 | 0.091 | 0.119 | 0.159 |
| EGF receptor signaling pathway | 0.031 | 0.080 | 0.134 | 0.178 | 0.240 |
| FAS signaling pathway | 0.180 | 0.328 | 0.311 | 0.508 | 0.689 |
| Gastrin_CCK2R_240212 | 0.106 | 0.232 | 0.382 | 0.459 | 0.642 |
| Hedgehog signaling pathway | 0.107 | 0.187 | 0.267 | 0.307 | 0.387 |
| Hypoxia response via HIF activation | 0.143 | 0.357 | 0.429 | 0.429 | 0.500 |
| Inflammation mediated by chemokine and cytokine signaling pathway | 0.046 | 0.131 | 0.210 | 0.282 | 0.389 |
| Insulin/IGF pathway-mitogen activated protein kinase kinase/MAP kinase cascade | 0.127 | 0.294 | 0.382 | 0.471 | 0.696 |
| Interferon-gamma signaling pathway | 0.086 | 0.229 | 0.305 | 0.324 | 0.457 |
| Interleukin signaling pathway | 0.045 | 0.116 | 0.170 | 0.229 | 0.306 |
| JAK/STAT signaling pathway | 0.173 | 0.320 | 0.360 | 0.493 | 0.680 |
| Notch signaling pathway | 0.125 | 0.299 | 0.326 | 0.403 | 0.604 |
| p38 MAPK pathway | 0.196 | 0.321 | 0.429 | 0.500 | 0.786 |
| PDGF signaling pathway | 0.045 | 0.111 | 0.185 | 0.231 | 0.305 |
| PI3 kinase pathway | 0.104 | 0.249 | 0.331 | 0.439 | 0.647 |
| T cell activation | 0.147 | 0.301 | 0.408 | 0.662 | 0.870 |
| TGF-beta signaling pathway | 0.049 | 0.128 | 0.200 | 0.278 | 0.360 |
| Toll receptor signaling pathway | 0.196 | 0.330 | 0.371 | 0.515 | 0.835 |
| VEGF signaling pathway | 0.094 | 0.164 | 0.228 | 0.292 | 0.363 |
| Wnt signaling pathway | 0.048 | 0.124 | 0.200 | 0.278 | 0.376 |

**Supplementary Table 5.** Pathway-interactome overlap for 22 PANTHER signaling pathways at interactome downsampling thresholds $t \in \{10, 30, 50, 70, 100\}$, where $t$ is the percentage of the interactome retained. Overlap is the fraction of a pathway's edges already present in the downsampled interactome, measured before the pathway edges are concatenated into the interactome. For each pathway and threshold, we resample the downsampled interactome up to 50 times, seeking a sample with at least 30 percent pathway overlap and full source-target connectivity in the combined network (every source can reach a target and every target is reachable from a source). We report the overlap of the first sample meeting both conditions, or the highest overlap across all attempts when none does.

#### 3 Coarse grid

The parameter ranges for each algorithm were constructed through the following process. First, we reviewed original publications and source code to identify documented defaults, recommended ranges, and any guidance on parameter behavior. The publications also helped us learn which parameters to prioritize for each algorithm, since some papers explain which parameters are important to tune. Parameters that were not included in grid search were set to their documented defaults, or excluded when no default was provided. Where the publication or software provided defaults and ranges, we treated these as starting points that reflect values the authors found effective, and built our grids around them. We added extreme values at both ends of each range to capture dataset-specific variation where the recommended ranges fail. Where needed, we spot-checked parameter ranges in the EGFR dataset, as described in the Pilot Study. For each parameter described below, further detail is available in the SPRAS documentation and, where applicable, the original manuscript for each algorithm.

**All Pairs Shortest Paths / BowTieBuilder:** No tunable parameters.

**DIAMOnD:** The manuscript recommend $n \approx 200$, reporting that the first 200 DIAMOnD genes participate in seed pathways at a rate similar to the seed proteins themselves and above random expectation.[3] The authors also find that $\alpha \approx 10$ performs best across datasets in the original publication.[3] We center the grids on these values and extend in both directions to capture dataset-specific variation.

**DOMINO:** The manuscript recommend a slice threshold of FDR $\leq 0.3$ for first-pass filtering and a module threshold of Bonferroni $\leq 0.05$ for final module selection.[4] Both grids are centered at the recommended values and extended slightly beyond to test cases where the defaults may not apply universally across datasets.

**Maximum Edge Orientation:** For the maximum path length parameter $k$, the manuscript used $k = 4$ and $k = 5$ in their primary evaluation and advise against $k \geq 6$, since the number of paths becomes computationally intractable.[5] The manuscript uses 20 iterations of random restarts[5]; we test values above and below to characterize the variance tradeoff. The `local_search` parameter is documented as a setting that should almost always be true, and setting it to false is strictly worse, so we set it to true for all runs.[5]

**MinCostFlow:** This method was adapted for pathway reconstruction, so its parameter grids were chosen empirically based on expected algorithm behavior. MinCostFlow output depends on the relationship between `flow` and `capacity`. When `capacity` > `flow`, the constraint is non-binding and the algorithm concentrates flow along the cheapest source-to-target paths, producing sparse, shortest-path-like output. When `capacity` < `flow`, the algorithm must split flow across multiple paths, producing denser output. The selected ranges span both cases. We used a simple dataset to spot-check candidate values, consolidating ranges that produced redundant output and extending them where the grid appeared too narrow. The final ranges are kept wide enough to allow for dataset-specific variation in the benchmarking study.

**NetMix2:** The manuscript describes setting the number of edges in the output modules to 40% of the original PPI network, rounded to the nearest 25,000.[9] The physical STRING interactome is 738,805 edges, so 40% corresponds to 295,522. We center the `num_edges` grid at this value with points spanning 5% to 50% of the network (specifically 5%, 15%, 25%, 35%, 40%, and 50% each rounded to the nearest 25,000). For the minimum edge density parameter `density`, we span the grid around the documented default of 0.05, with values chosen empirically to capture dataset-specific variation.

**Omics Integrator 1:** Following the identification of $\omega$, $\beta$, and $\mu$ as the primary solution-determining parameters in the manuscript, we focus on tuning these along with `D`. For each parameter, we grid around the ranges recommended in the paper[10]: $\mu \in [0.0001, 0.1]$, $\beta \in [1, 20]$, `D` $\in [5, 15]$, and $\omega \in [1, 10]$. Grids were selected by spot-checking on an simple dataset, using the published defaults and recommended ranges as initial references and testing whether the chosen values produced feasible runs. We restricted tuning to these four parameters since Omics Integrator 1 is computationally expensive, and expanding the grid to additional parameters or finer spacing would produce combination counts beyond what is feasible to run across the benchmarking study.

Several Omics Integrator 1 parameters were excluded from the grid. The ensembling parameters (`noisy_edges`, `shuffled_prizes`, `random_terminals`, and parameter `r`) were left at their defaults as SPRAS has its own ensembling and does not directly support the type of ensembling these parameters control. The prize-modification flags `mu_squared` and `exclude_terms` were also fixed at their defaults to limit grid size. Parameter `g` was fixed at its default of 0.001, since it primarily controls the convergence criteria rather than the type of optimal solution.

**Omics Integrator 2:** No paper or documented parameter ranges accompany Omics Integrator 2's defaults of $\omega = 6$, $\beta = 1$, and $\gamma = 20$.[11] Example configurations in the Omics Integrator 2 repository use small grids spanning $\omega \in [0.25, 1]$, $\beta \in [0.25, 2]$, and $\gamma \in [3, 4.5]$, which provide a starting reference for parameter behavior.[11] Grids for $\omega$, $\beta$, and $\gamma$ were selected through spot-checking on a simple dataset, using the published defaults and example configurations as initial references and testing on the dataset to identify extremes.

Although Omics Integrator 1 and Omics Integrator 2 share $\omega$ and $\beta$, the two tools suggest scaling these parameters differently (e.g., Omics Integrator 1 suggests $\beta \in [1, 20]$ while Omics Integrator 2's suggest $\beta \in [0.25, 2]$), so we did not reuse Omics Integrator 1's ranges. Note also that both tools expose a parameter named $\gamma$, but it plays a different role in each: in Omics Integrator 1 it is the msgsteiner reinforcement parameter, whereas in Omics Integrator 2 it is the degree-based edge penalty.

The ensembling parameters `noisy_edges`, `random_terminals`, and `noise` were left at their defaults, since SPRAS does not currently support the re-run aggregation these parameters are intended to drive.

**PathLinker:** The manuscript evaluates up to $k = 20000$ but reports that most pathway proteins appear within the first $\sim 150$ paths.[12] We grid more densely in the $k = 10$ to $k = 300$ range and include $k = 1$, $k = 1000$, $k = 5000$ and $k = 20000$ as extreme points to capture any dataset-specific variation.

**Random Walks with Restarts / Source-Target Random Walks with Restarts:** The PageRank manuscript proposes an $\alpha$ value of 0.85.[17] Our grid spans values from restart-dominated behavior, where the walker stays local to sources, to walk-dominated behavior, where the walker spreads broadly across the network. For `threshold`, the number of nodes to retain in the returned subgraph, subgraph sizes range from 50 to 500, chosen empirically

**ResponseNet:** The manuscript reports that ResponseNet has a narrow effective range for $\gamma$: values below 7 typically yield empty solutions, values above 20 saturate the maximum attainable flow, and $\gamma = 10$ produces intermediate-sized networks. The authors also note that changes of $\pm 1$ rarely affect the top-ranked proteins.[15] The grid spans the reported range, with extremes $\gamma = 5$ and $\gamma = 25$ to capture any dataset-specific variation.

**TieDIE:** The TieDIE manuscript and accompanying code document only the alpha parameter, providing no guidance on the remaining parameters or their defaults.[16] Parameter values were instead chosen close to the defaults where these could be inferred. The default for $c$ is 3, so values from 3 to a maximum of 7 were chosen. The default for $s$ is 1, so values of 1, 3, and 5 were chosen. The `pagerank` flag, which enables personalized PageRank diffusion, and the `all_paths` flag, which uses all paths rather than only causal paths, we set as True, as we don't provide signed edges.

| Algorithm | Coarse Grid |
|---|---|
| All Pairs Shortest Paths | None |
| BowTieBuilder | None |
| DIAMOnD | `n`: [10, 50, 100, 150, 200, 500, 1000]<br>$\alpha$: [1, 3, 5, 10, 15, 25, 50] |
| DOMINO | `slice_threshold`: [0.05, 0.1, 0.3, 0.5]<br>`module_threshold`: [0.005, 0.01, 0.05, 0.1] |
| Maximum Edge Orientation | `max_path_length`: [2, 3, 4, 5]<br>`local_search`: [True]<br>`rand_restarts`: [5, 20, 50] |
| MinCostFlow | `flow`: [1, 5, 10, 25, 50, 100]<br>`capacity`: [1, 5, 10, 100] |
| NetMix2 | `num_edges`: [25000, 100000, 175000, 250000, 300000, 375000]<br>`density`: [0.02, 0.05, 0.1, 0.2, 0.35, 0.5] |
| Omics Integrator 1 | $\omega$: [0.5, 1, 5, 10]<br>$\beta$: [1, 5, 10, 15, 20]<br>`D`: [5, 10, 15]<br>$\mu$: [0.0, 0.0001, 0.005, 0.01] |
| Omics Integrator 2 | $\omega$: [0.25, 0.5, 1, 3, 6, 15]<br>$\beta$: [0.25, 0.75, 1, 5, 10, 15]<br>$\gamma$: [0, 1, 3, 4, 5, 20] |
| PathLinker | `k`: [1, 10, 50, 100, 200, 500, 1000, 5000, 20000] |
| Random Walks with Restarts | $\alpha$: [0.05, 0.3, 0.5, 0.7, 0.85, 0.99]<br>`threshold`: [50, 100, 300, 500] |
| ResponseNet | $\gamma$: [5, 7, 8, 10, 15, 19, 20, 25] |
| Source-Target Random Walks with Restarts | $\alpha$: [0.05, 0.3, 0.5, 0.7, 0.85, 0.99]<br>`threshold`: [50, 100, 300, 500] |
| TieDIE | `c`: [3, 7]<br>`s`: [1, 3, 5]<br>`pagerank`: [True, False]<br>`all_paths`: [True] |

**Supplementary Table 6.** Coarse parameter grids for pathway reconstruction algorithms. Total number of parameter combinations is 672.

#### 4 Heuristic survey

To select topological heuristics and their ranges for two-stage parameter tuning, a survey will be distributed to SPRAS team members. The survey will elicit intuitions about the structural properties of a plausible reconstructed pathways.[18] Respondents will be given the following framing: we run a pathway reconstruction algorithm across a parameter grid, and graph properties are needed to filter out implausible outputs such as a single edge, a single node, or a fully connected graph.

The survey will collect respondent intuitions along several structural dimensions, each corresponding to a pathway summary statistic defined in the evaluation metrics. For each statistic, respondents will complete four fields: a minimum value, a maximum value, an ideal value or range, and their reasoning. Respondents can also use the reasoning field to indicate that a given statistic was not useful or provide other context. These responses will determine which graph topological heuristics are used and what ranges are set for two-stage parameter tuning.

#### 5 SPRAS File Formats

**Node Files** Node files contain information about the input nodes specifying which nodes are prizes, sources, targets, active status, or dummy nodes. Node files contain a header line.

| NODEID | prize | sources | targets | active | dummy |
|---|---|---|---|---|---|
| A | 1.0 | | True | True | True |
| B | 3.3 | True | | True | |
| C | 2.5 | | True | True | |
| D | 1.9 | True | True | True | |

**Supplementary Table 7.** Example node file format

**Edge Files** Edge files represent the background interactomes. Each row specifies two connected nodes, an edge weight, and optionally a directionality indicator. If the directionality column is absent, SPRAS assumes all edges are undirected.

| | | | |
|---|---|---|---|
| A | B | 0.98 | U |
| B | C | 0.77 | D |
| C | D | 0.65 | U |

**Supplementary Table 8.** Example edge file with directionality column.

| | | |
|---|---|---|
| A | B | 0.98 |
| B | C | 0.77 |
| C | D | 0.65 |

**Supplementary Table 9.** Example edge file without directionality column.

**Gold Standard Files** Gold standard files provide reference sets of nodes or edges for evaluating reconstruction accuracy. Unlike node and edge input files, gold standards are not converted for individual algorithms but remain in their standardized format for post analysis evaluation purposes.

| |
|---|
| A |
| B |
| C |
| D |

**Supplementary Table 10.** Example gold standard node file. The file contains one node identifier per line.

| | |
|---|---|
| A | B |
| B | C |
| C | D |

**Supplementary Table 11.** Example gold standard edge file. The file contains two columns, with each row representing an edge between two nodes.

| | | |
|---|---|---|
| A | B | U |
| B | C | D |
| C | D | U |

**Supplementary Table 12.** Example gold standard edge file. The file contains two columns, with each row representing an edge between two nodes with directionality included.

**Output Files** Our output files take the raw pathways generated by each algorithm and standardize them into the SPRAS output file format. Each row represents an edge, given as two connected nodes, a rank for that edge, and a directionality indicator. Rank is currently only meaningful for PathLinker and DIAMOnD. All other algorithms default to a constant rank of 1. The output files contain a header line.

| Node1 | Node2 | Rank | Direction |
|---|---|---|---|
| A | B | 1 | U |
| B | C | 1 | D |
| C | D | 1 | U |

**Supplementary Table 13.** Example standardized output pathway format.

#### 6 Computational Performance Assessment

| Pathways Chosen for Computational Performance Assessment | Used in Input Node Experiment | Used in Interactome Experiment |
|---|---|---|
| Balanced pathway | ✓ | |
| Cadherin signaling pathway | ✓ | ✓ |
| Gastrin_CCK2R_240212 | ✓ | ✓ |
| Hedgehog signaling pathway | ✓ | ✓ |
| Inflammation mediated by chemokine and cytokine signaling pathway | ✓ | ✓ |
| Interleukin signaling pathway | ✓ | ✓ |
| JAK/STAT signaling pathway | ✓ | ✓ |
| Mega pathway | ✓ | |
| Notch signaling pathway | ✓ | ✓ |
| PDGF signaling pathway | ✓ | ✓ |
| p38 MAPK pathway | ✓ | ✓ |
| Source-focused pathway | ✓ | |
| Target-focused pathway | ✓ | |
| TGF-beta signaling pathway | ✓ | ✓ |
| Wnt signaling pathway | ✓ | ✓ |

**Supplementary Table 14.** We will use 15 pathways (11 PANTHER signaling pathways and the four merged variant pathways) for computational performance assessment. The four merged variant pathways are excluded for the interactome experiment.

### 7 Running at Scale

| Algorithm | Parameter combinations | | | Jobs by dataset | | | | |
|---|---|---|---|---|---|---|---|---|
| | Coarse grid | Fine grid | Worst case | DepMap | PANTHER | EGFR | DISEASES | Total jobs |
| All Pairs Shortest Paths | 1 | 0 | 1 | 627 | 70 | 1 | 0 | 698 |
| BowTieBuilder | 1 | 0 | 1 | 627 | 70 | 1 | 0 | 698 |
| DIAMOnD | 49 | 120 | 169 | 105,963 | 11,830 | 169 | 20,956 | 138,918 |
| DOMINO | 16 | 33 | 49 | 30,723 | 3,430 | 49 | 6,076 | 40,278 |
| NetMix2 | 36 | 85 | 121 | 75,867 | 8,470 | 121 | 15,004 | 99,462 |
| Omics Integrator 1 | 240 | 1,965 | 2,205 | 1,382,535 | 154,350 | 2,205 | 273,420 | 1,812,510 |
| Omics Integrator 2 | 216 | 1,115 | 1,331 | 834,537 | 93,170 | 1,331 | 165,044 | 1,094,082 |
| Random Walks with Restarts | 24 | 53 | 77 | 48,279 | 5,390 | 77 | 9,548 | 63,294 |
| Maximum Edge Orientation | 12 | 23 | 35 | 21,945 | 2,450 | 35 | 0 | 24,430 |
| MinCostFlow | 24 | 53 | 77 | 48,279 | 5,390 | 77 | 0 | 53,746 |
| PathLinker | 9 | 8 | 17 | 10,659 | 1,190 | 17 | 0 | 11,866 |
| ResponseNet | 8 | 7 | 15 | 9,405 | 1,050 | 15 | 0 | 10,470 |
| Source-Target Random Walks with Restarts | 24 | 53 | 77 | 48,279 | 5,390 | 77 | 0 | 53,746 |
| TieDIE | 12 | 18 | 30 | 18,810 | 2,100 | 30 | 0 | 20,940 |
| Total | 672 | 3,533 | 4,205 | 2,636,535 | 294,350 | 4,205 | 490,048 | 3,425,138 |

**Supplementary Table 15.** Parameter combinations and total reconstruction jobs by algorithm and dataset. DepMap, PANTHER, EGFR, and DISEASES comprise 627, 70, 1, and 124 dataset instances respectively. The worst-case parameter combination cdount is the sum of the coarse and fine grid combinations, the maximum number of parameter settings run per dataset instance. Jobs per dataset equal this worst-case count multiplied by the instance count. DISEASES runs only the six algorithms that use prizes or active nodes, so the other algorithms do not contribute to the number of DISEASES jobs.

| Step | Jobs |
|---|---|
| `all` | 1 |
| `log_datasets` | 822 |
| `log_parameters` | 4,205 |
| `merge_input` | 822 |
| `merge_gs` | 822 |
| `parse_output` | 3,425,138 |
| `prepare_input` | 10,516 |
| `reconstruct` | 3,425,138 |
| Total | 6,867,464 |

**Supplementary Table 16.** Total job counts at worst-case total parameter combinations. `log_datasets`, `merge_input`, and `merge_gs` run once per dataset instance (822 instances). `prepare_input` runs once per dataset instance per algorithm: the 698 non-DISEASES instances run on all 14 algorithms and the 124 DISEASES instances run on 6, giving $698 \times 14 + 124 \times 6 = 10{,}516$. `log_parameters` uses the worst-case total parameter combinations summed across all algorithms. `parse_output` and `reconstruct` each equal the total reconstruction job count, since every reconstructed pathway in the worst case must also be parsed.
